# Systematic single-cell analysis reveals dynamic control of transposable element activity orchestrating the endothelial-to-hematopoietic transition

**DOI:** 10.1101/2023.06.19.545461

**Authors:** Cong Feng, Ruxiu Tie, Saige Xin, Yuhao Chen, Sida Li, Xiaotian Hu, Yincong Zhou, Yongjing Liu, Yueming Hu, Yanshi Hu, Hang Pan, Zexu Wu, Haoyu Chao, Shilong Zhang, Qingyang Ni, Jinyan Huang, Wenda Luo, He Huang, Ming Chen

**Affiliations:** Department of Bioinformatics, Zhejiang University College of Life Sciences, Hangzhou 310058, China; Bioinformatics Center, The First Affiliated Hospital, Zhejiang University School of Medicine, Hangzhou 310058, China; Bone Marrow Transplantation Center, the First Affiliated Hospital, Zhejiang University School of Medicine, Hangzhou 310058, China; Liangzhu Laboratory, Zhejiang University Medical Center, Hangzhou 310058, China; Institute of Hematology, Zhejiang University, Hangzhou 310058, China; Zhejiang Province Engineering Laboratory for Stem Cell and Immunity Therapy, Hangzhou 310058, China; Department of Hematology, The Second Clinical Medical College of Shanxi Medical University, Shanxi Medical University, Taiyuan 030000, China; Department of Hematology-Oncology, Taizhou Hospital of Zhejiang Province, Linhai 317000, China; Department of Veterinary Medicine, Zhejiang University College of Animal Sciences, Hangzhou 310058, China

**Keywords:** Endothelial-to-hematopoietic transition, Transposable element, Hematopoietic stem cell, Inflammatory signaling, Cis-regulatory element, Hypoxia

## Abstract

**Background:** The endothelial-to-hematopoietic transition (EHT) process during definitive hematopoiesis in vertebrate is highly conserved. Stage-specific expression of transposable elements (TEs) has been detected during zebrafish EHT and may promote hematopoietic stem cell formation by activating inflammatory signaling. However, little is known about how TEs contribute to the EHT process in human and mouse.

**Results:** We reconstructed the single-cell EHT trajectories of human and mouse, and resolved the dynamic expression patterns of TEs during EHT. Most TEs presented a transient co-upregulation pattern along the conserved EHT trajectories. Enhanced TE activation was tightly associated with the temporal relaxation of epigenetic silencing systems. TE products can be sensed by multiple pattern recognition receptors, triggering inflammatory signaling to facilitate the emergence of hematopoietic stem cells. Furthermore, we observed that hypoxia-related signals were enriched in cells with higher TE expression. Additionally, we constructed the hematopoietic cis-regulatory network of accessible TEs and identified potential enhancers derived by TEs, which may boost the expression of specific EHT marker genes.

**Conclusions:** Our study provides a systematic vision on how TEs are dynamically controlled to promote the hematopoietic fate decision through transcriptional and cis-regulatory networks, and pre-train the immunity of nascent hematopoietic stem cells.

## Introduction

Hematopoietic stem cells (HSCs) are a pluripotent cell population in the blood system, which possess the ability of self-renewal and lineage differentiation to maintain the function of the hematopoietic system throughout the lifespan. In embryos, HSCs originate from a subpopulation of endothelial cells (ECs) with hematopoietic potential in the aorta-gonad-mesonephro (AGM) region, which have been directly tracked by using time-lapse imaging methods [1-5]. The endothelial-to-hematopoietic transition (EHT) process is highly conserved in vertebrate embryos including zebrafish, mouse, and human [6]. HSCs first emerge during embryonic day (E) 10.5-11.5 in mouse AGM [7, 8] and Carnegie stage 13-17 (CS13-17; 4-6 weeks) in human AGM [9, 10], and migrate to the fetal liver under the promotion of blood flow, and finally colonize the fetal bone marrow. The identification and functional analysis of heterogeneous and intermediate cell clusters during EHT *in vivo* has been a challenging task in understanding and probing embryonic hematopoiesis [11]. It is presented that a specific cluster of arterial endothelial cells (AECs) from primitive blood vessels undergoes hemogenic fate decision to become HSC-primed hematopoietic endothelial cells (HECs) [12-14]. In addition to HECs, at least two kinds of HSC precursors (distinguished by CD45) are found in mouse intra-aortic hematopoietic clusters (IAHCs) [15-18]. In the last years, the rapid development of single-cell sequencing technologies has greatly broadened the insights into cellular heterogeneity and complex relationships during developmental hematopoiesis [19, 20]. Using single-cell RNA sequencing (scRNA-seq) and single-cell sequencing assay for transposase-accessible chromatin (scATAC-seq), the continuous EHT trajectory has been constructed and an intermediate cell population proximal to HECs is identified (termed pre-HECs) in mouse [21, 22]. Recently, a research team discovered a signature gene set RUNX1+HOXA9+MLLT3+MECOM+HLF+SPINK2+ that distinguishes human HSCs from other hematopoietic progenitor cells, and for the first time depicted the single cell landscape of HSCs from origination to maturation in the human embryo [23]. However, whether the cell types are comparable in human and mouse EHT, and how many marker genes are conserved, has not been systematically investigated.

It is recognized that the EHT process is strictly regulated by multiple regulatory factors at the transcriptional and epigenetic levels [24, 25]. Various transcription factors such as Runx1, Gfi1 and Gata2 have been shown to play vital roles in HSC development [26-28]. Signaling pathways like NOTCH, WNT, YAP and VEGF are also involved in HSC fate decision [25]. Besides, it is worth noting that some inflammatory signals are highlighted to regulate the HSC emergence [29, 30]. For example, interleukins like IL-1, IL-3, and IL-6 regulate HSC development endogenously in the AGM region [31, 32]. Tumor necrosis factors (TNF) and interferon signals (IFN) can also promote the development of HSCs [33, 34]. In the innate immune system, pattern recognition receptors (PRRs) such as Toll-like receptors (TLRs), RIG-I-like receptors (RLRs), NOD-like receptors (NLRs), and C-type lectin receptors (CLRs) are considered to be key activators of these inflammatory responses [35]. It has been pointed out that Toll-like receptor 4 (TLR4) can regulate the formation and development of HSCs by promoting Notch activity through MyD88-mediated NF-κB signaling [30]. A recent study in zebrafish unraveled that RLRs (including RIG-I, MDA5 and LGP2) are also involved in HSC formation through activating downstream inflammatory signaling such as TNF receptor-associated factors (TRAFs) [36]. Typically, these PRRs induce antiviral immune responses by recognizing single-stranded RNAs (ssRNAs) or double-stranded RNAs (dsRNAs) produced by exogenous pathogens [37, 38]. During the formation of HSCs, the ligands for PRRs are puzzling because the AGM region is supposed to be in a sterile niche. Nonetheless, transposable elements (TEs) abundantly distributed throughout the genome may provide endogenous nucleic acids for PRRs [39]. Strikingly, the expression TEs has been detected during zebrafish EHT, and is demonstrated to affect HSC generation through the RLR pathway [36]. However, as yet, little is known about the contributions of TEs to human and mouse EHT.

TEs are consisting of retrotransposons and DNA transposons (DNAs). Retrotransposons transpose by a copy-and-paste mechanism, whereby an RNA intermediate is reverse-transcribed and then inserted into a new genomic locus. Most of the TEs in human and mouse are retrotransposons, whether long interspersed nuclear elements (LINEs), short interspersed nuclear elements (SINEs) or hominid SVAs (SINE-VNTR-Alu), and long terminal repeats (LTRs), most of which also known as endogenous retroviruses (ERVs). The jumping mechanism of TEs may induce genome instability, and uncontrolled transposition can lead to disease [40, 41]. Therefore, a variety of defense systems have evolved to domesticate TEs, including chromatin modification, small RNA silencing, and post-transcriptional repression [42, 43]. In vertebrates, Krüppel-associated box zinc finger protein (KRAB-ZFP) is one of the prominent TE silencing systems, which inhibits TEs through interacting with KAP1 to recruit DNA methyltransferases (DNMT), SETDB1, HP1 and the nucleosome remodeling deacetylase (NuRD) complex [42, 44]. The human silencing hub (HUSH) complex coupled with the ATPase MORC2 to deposit H3K9me3 for de novo silencing of TEs. Although the activity of TEs in the genome is often silenced, they can be activated in a temporary or tightly fashion both at transcriptional and epigenetic levels to shape embryonic development [45, 46]. Studies at the single cell level show that TEs have cell type-specific expression during gastrulation and organogenesis, and participate in dynamic regulation of pluripotency reprogramming and lineage differentiation [47-49]. There are also related evidences show that TEs can contribute to the hematopoietic regeneration and fate decision [50-52], which may provide new insights on therapies of certain hematopoietic diseases.

Despite progress in understanding the expression and potential RLR pathway of TEs during zebrafish EHT, there is still limited exploration of the single-cell expression and regulatory landscape of TEs, as well as the underlying mechanisms of TE activation in human and mouse. In this study, we first performed a comprehensive survey of the genomic landscape and potential regulatory functions of TEs in human and mouse, laying a foundation for further analysis of TE activity in embryonic HSC development. We reconstructed the human and mouse EHT trajectories using scRNA-seq datasets, demonstrated the conservation of EHT cell types between species, and resolved the dynamic expression patterns of TEs during EHT. We observed that most TEs presented low cell type-specificity on the conserved EHT trajectories, but form a co-upregulation pattern during pre-HEC specification. The activation of TEs in the pre-HEC stage was strongly associated with the transient relaxation of several TE silencing systems. The delayed TE product sensing through various PRRs can trigger inflammatory signaling to facilitate immune activation of HSCs. Importantly, these observations are highly conserved in human and mouse. Additionally, by analyzing scATAC-seq data, we constructed the hematopoietic cis-regulatory network of accessible TEs and identified two potential TE derived enhancers, which may promote the expression of a specific pre-HEC marker (Gja5). Furthermore, by combining the spatial transcriptome of human AGM and a bulk RNA-seq of hypoxia response, we hypothesized that the hypoxic AGM niche may be partially responsible for transient TE activation preceding hematopoietic fate commitment. Taken together, our study provides a systematic single-cell investigation into the contribution of TEs to the expression and regulatory landscape of EHT, which may shed lights on studying TEs in the context of stem cell development and other cell type transition systems.

## Results

### Widespread TEs harbor great regulatory potential in human and mouse

TEs in human and mouse genomes can be divided into 4 classes (LINEs, SINEs, LTRs and DNAs), and the hominids SVAs (6 families) are included into SINEs for calculation. These TEs can be further classified into 42 superfamilies (1176 families) and 41 superfamilies (1256 families) in human and mouse, respectively (Additional file 1: Table S1 and S2). Collectively, TEs account for about 46.38% and 41.76% of the human and mouse genomes, respectively (Fig. 1A, E; Additional file 1: Table S3 and S4). The majority of TEs are located in non-coding regions, including intergenic regions, introns and UTRs (Fig. 1B, F; Additional file 1: Table S5 and S6). Interestingly, TEs are less distributed near the transcription start sites (TSS) and transcription termination sites (TTS) (Fig. 1C, G), which may be important for maintaining the specificity of gene transcription [53]. To explore the regulatory potential of TEs, we calculated the copy number of each TE superfamily overlapping with CpG islands and candidate cis-regulatory elements (cCREs, downloaded from ENCODE-SCREEN [54]). In the human genome, SINEs/SVAs (including Alu and SVA) contribute more than 39% of CpG islands (Fig. 1D; Additional file 2: Table S1 and S13), but in mouse, SINEs only overlap with about 1.47% of CpG islands, while LINEs (especially L1) and LTRs (ERV1 and ERVK) both contribute more than 15% of CpG islands (Fig. 1H; Additional file 2: Table S2 and S14). In most cases, abundant CpG sites keep TEs repressed in the methylated state. However, through demethylation processes such as epigenetic reprogramming, it is possible for the TEs to be activated and play a role in embryonic development [55, 56]. Among the cCREs, in addition to promoter-like sites (PLS), a considerable proportion (36.39%-57.55% in human and 21.34%-42.95% in mouse) of proximal enhancer-like signatures (pELS), distal enhancer-like signatures (dELS), CTCF signatures and DNase-H3K4me3 signatures have intersections with TEs (Figure 1D, 1H; Additional file 2: Table S3-S12). These evidences suggest that widely distributed TEs have evolved enormous regulatory potential and may exert unique contributions in pluripotency and early embryogenesis both at transcriptional and epigenetic levels [57-61]. In this study, we will focus on analyzing the expression and chromatin accessibility of TEs during EHT, expecting to reveal the potential regulatory mechanism of TE in the formation of HSCs.

**Figure 1.**
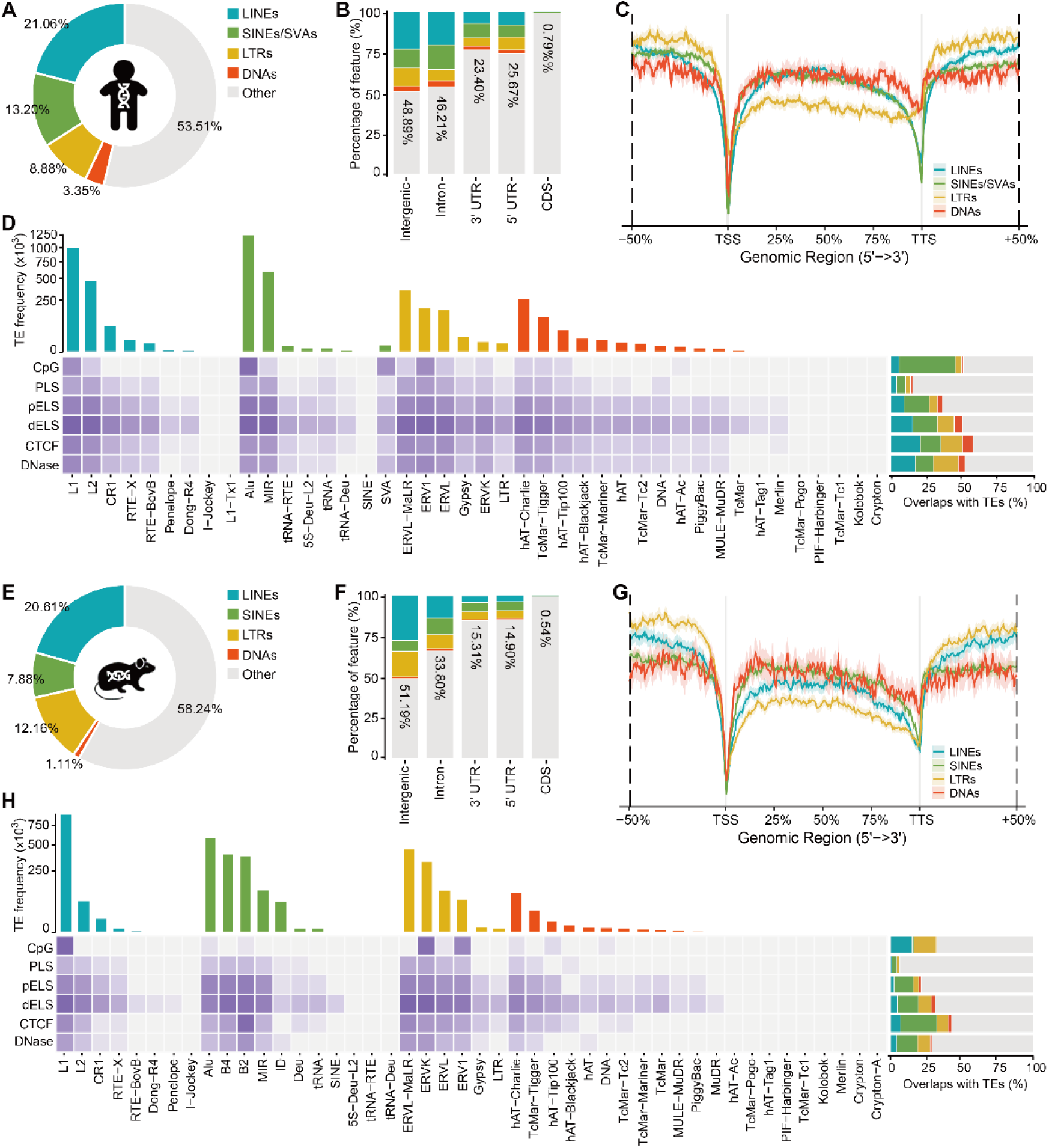
Genomic landscape of TEs in human and mouse. **A, E** Genome coverage of TEs in human and mouse. **B, F** Overlaps of TEs with gene structures in human and mouse. **C, G** Distribution of TEs along the gene body in human and mouse. **D, H** The frequency of TE superfamilies (upper bar plot) and overlaps of TEs with CpG islands and cis-regulatory elements (heatmap and right bar plot) in human and mouse.

### TEs are detected with low cell type-specificity on the conserved EHT trajectories of human and mouse

Accurate cell subtyping is the basis for studying various molecular regulatory activities during EHT. With the aid of published scRNA-seq data of the human and mouse AGM (Additional file 3: Table S1) [22, 23], we reconstructed the EHT trajectories *de novo* by analyzing RNA velocities (Fig. 2A, E). Interestingly, we observed aberrant orientation of RNA velocity during the progression from pre-HECs to HSCs (which was also reflected in the latent time) in both human and mouse, suggesting a differentiation bottleneck during hematopoietic specification [22]. The cell type annotation of the AGM region and the characterization of EHT cell subtypes were relied on the known marker genes and signatures proposed in the human EHT study [23]. First, we isolated cell clusters of ECs and HSCs from the AGM UMAP, and obtained EHT clusters through dimension reduction and clustering. By using common marker genes of VECs (CDH5, NRP2, NR2F2), AECs (GJA4, HEY1, DLL4), pre-HECs (TMEM100, GJA5, EDN1) and HSCs (RUNX1, MYB, HLF), we accurately identified these four cell types within the EHT cell clusters (Fig. 2B, F; Additional file 4: Fig. S1A-C, F and Fig. S2A-C, F). However, distinguishing HECs directly from the cluster level can be challenging since they serve as transitional cells from pre- HECs to HSCs. Therefore, HECs were selected based on the co-expression of CDH5, RUNX1 and MYCN, along with the absence of PTPRC (Fig. 2B, F; Additional file 4: Fig. S1D-F and Fig. S2D-F). After reconstructing the EHT trajectory, we integrated the human and mouse EHT data based on the shared homologous genes. The results showed that the EHT of both showed a highly conserved pattern, although a relatively larger number of HECs were captured in mouse (Additional file 4: Fig. S3A, B). The majority of EHT marker genes were found to be conserved in human and mouse. For instance, ACE is positive, CD44 is low and KIT is negative in pre-HEC [21, 62]. However, there are also some species-specific EHT marker genes, such as IL33 and SPINK2 only express in human pre-HECs and HSCs, respectively, while in mouse, Ikzf2 is more enriched HECs and HSCs (Additional file 4: Fig. S3D).

**Figure 2.**
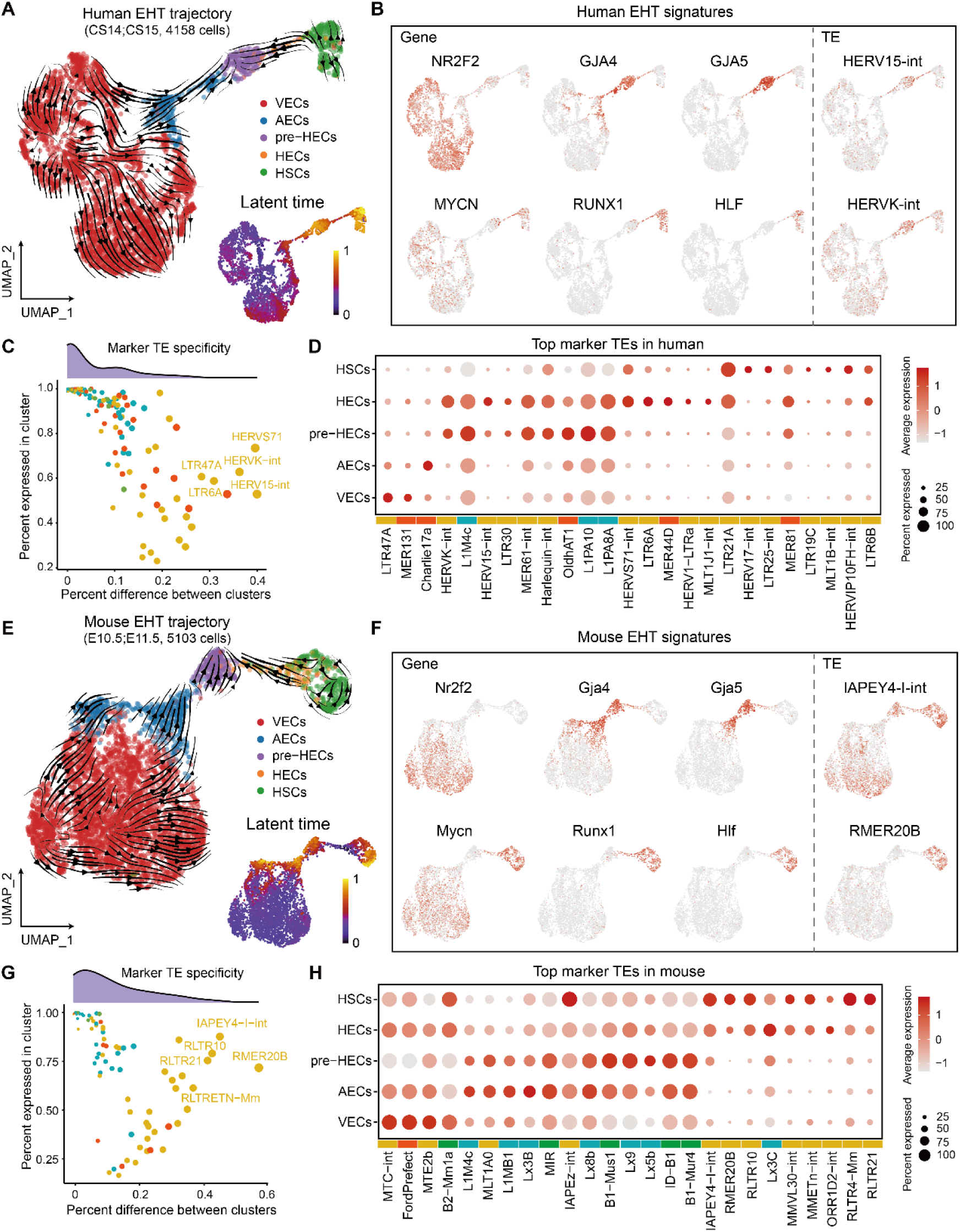
The EHT trajectories and TE expression overview in human and mouse. **A, E** Human and mouse EHT trajectories and predicted latent time. **B, F** Human and mouse EHT signatures. NR2F2 marks VECs, GJA4 marks AECs, GJA5 marks late AECs and pre- HECs, MYCN and RUNX1 marks HECs and HSCs, HLF marks HSCs. These markers are conserved between human and mouse. Only a few marker TEs were identified in the analysis. **C, G** The specificities of marker TEs in human and mouse. The majority of TEs with high cell type-specificities are LTRs. **D, H** Top markers of each cell type in human and mouse EHT.

To explore the dynamics of TEs on the EHT trajectories, we computed the family-level TE expression to each EHT cell. Through differential expression analysis, we identified 214 and 96 marker TEs in human and mouse EHT cells (average log2FC≥0.25 and adjusted P-value≤0.05), respectively. Among these, 198 TEs (92.52%) in human were enriched in pre-HECs, while 72 TEs (75%) in mouse were belonged to AECs and pre-HECs (Additional file 5: Table S1 and Table S2). Notably, in human EHT, HERV15-int and HERVK-int appeared to be enriched in pre-HECs and HECs, whereas in mouse EHT, IAPEY4-I-int and RMER20B were highly expressed in HSCs (Fig. 2B, F). It should be noted that the specificities of most marker TEs are not significant, that is, although TEs are enriched in the target cell population, the differences in expression percentage compared with other cell populations are mostly less than 0.2 (Fig. 2C, G). The top marker TEs for each cell type of human and mouse EHT were displayed in Fig. 2D, H. Surprisingly, the marker TEs that showed relatively higher specificities (percent difference >0.25) are mostly ERVs, which is consistent in both human and mouse. In particular, among the relatively highly specific marker TEs in human EHT (Fig. 2C), primate-restricted HERVK transcripts have been reported to be abundant in primordial germ cells, endodermal cells and blood progenitors during human gastrulation, whereas HERVS71 transcripts are frequently detected in primitive streak, nascent mesoderm and definitive endoderm [48]. These observations provide valuable insights into the regulatory roles of TEs to EHT and early embryogenesis.

### TEs form a distinguished co-upregulation pattern during pre-HEC specification

To identify modules that may participate in common regulatory processes during EHT, we clustered the dynamic expression profiles of genes and TEs by co-expression network analysis. A total of 988 filtered TEs, 528 marker genes (average log2FC≥0.5 and adjusted P-value≤0.05) in human and 864 filtered TEs, 421 marker genes in mouse were included for co-expression analysis (Additional file 5: Table S3 and S4). Those selected genes and TEs were clustered into 5 modules (HME1-5 and MME1-5) in both human and mouse, in which HME1-4 (MME1-4) were enriched in VECs, AECs, pre-HECs and HECs/HSCs, respectively (Fig. 3A-D; Additional file 6: Table S1 and S2). The top 10 hub genes/TEs of each module were listed in Fig. 3B and Fig. 3D, characterizing the features that may play a leading role in each module. Some conserved hub genes can be found in human and mouse, such as GJA5 and TMEM100 are hub genes in pre-HECs, while MYB, SPI1 and CORO1A are hub genes in HSCs. Interestingly, most TEs (645 of 670 in human and 659 of 699 in mouse) were clustered in HME5 and MME5 (Fig. 3B, D). The expression patterns of these TEs were that, there was a consistent expression in almost the whole EHT process, but specifically showed an upward trend in pre-HECs (appeared earlier in the AEC stage in mouse). Among the TEs included in HME5 and MME5, although LTRs accounted for the largest proportion (39.84% and 56.90% in human and mouse, respectively), their module connectivity scores (kME scores) were relatively low (Fig. 3E, F). In contrast, SINEs and LINEs exhibited higher kME scores in both HME5 and MME5. It is evidenced that overexpression of a SINE copy (sine3-1a) can enhance HSC formation in zebrafish [36]. In particular, among TEs with kME scores greater than 0.3, L1 was the most abundant in both human and mouse (94 in 190 and 68 in 179) (Additional file 6: Table S1 and S2). This observation could be attributed to the abundant CpG islands on the L1 elements, implying a potential reduction in methylation levels of TEs during the pre-HEC stage. In addition, 260 common TEs in human and mouse are found to be enriched in HME5 and MME5 (most with the top average kME scores are L1 elements) (Additional file 6: Table S3). Some top-ranked (according to kME scores) common TEs (L1, L2 and MIR) and species-specific TEs (Alu and mouse-specific B2 and B4) were displayed in Fig. 3G, H.

**Figure 3.**
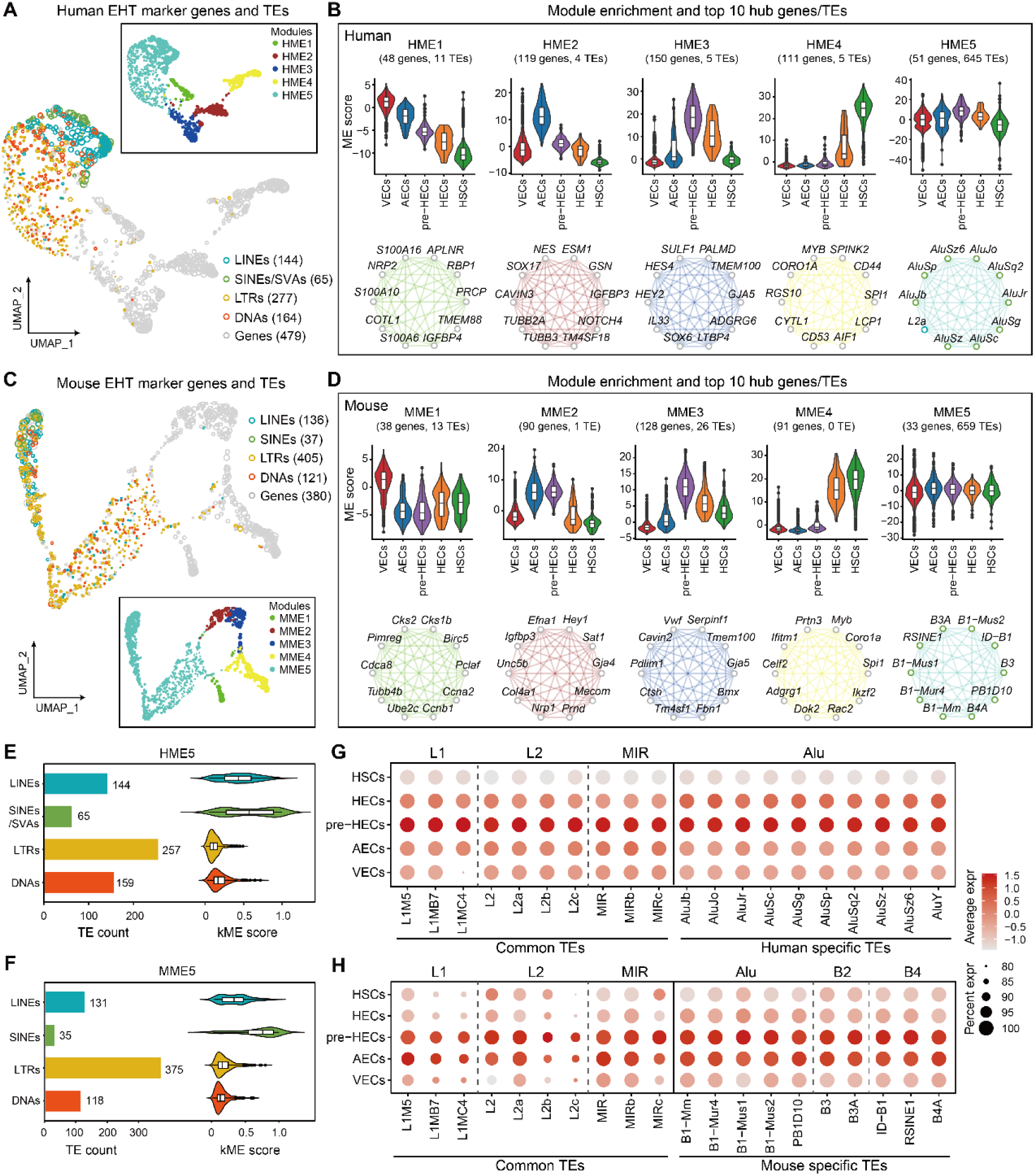
Co-expression network analysis of marker genes and expressed TEs. **A, C** Co-expression modules of marker genes and TEs in human and mouse. Most TEs tend to cluster together as distinct modules (HME5 and MME5). **B, D** The expression patterns and top 10 hub genes/TEs of each module in human and mouse. TEs show a common upregulation trend in pre-HECs. In the case of mouse, this upregulation appears to occur even earlier, during the AEC stage. **E, F** TE composition (bar plot) and module connectivity (kME, violin plot) of HME5 and MME5. **G, H** Dot plots show the expression levels of selected common and species-specific TEs.

### TE silencers are transiently relaxed in pre-HECs

We performed differential expression analysis on pre-HECs against VECs and HSCs, and the results showed that TEs were pervasively upregulated in pre-HECs in human (Fig. 4A; Additional file 7: Table S1 and S3), while this phenomenon was relatively insignificant in mouse (Fig. 4B), but the upregulated TEs in pre-HECs were still more than other cell types (Additional file 7: Table S5 and S7). This is consistent with our observation of TE-enriched modules (HME5 and MME5) upregulated at the pre-HEC stage in co-expression network analysis (Fig. 3B, D). However, in mouse, it seems that there is already a large-scale TE activation in AECs, which may be related to the presence of more pre-HEC-primed cells in AECs (Fig. 3D). It can also be seen that the fold changes of differentially expressed TEs were generally low, which also explained why we identified few cell type-specific marker TEs during EHT (Fig. 2C, G). Since TEs are normally repressed in most cases, it is reasonable to assume that the transient activation of TEs in pre-HECs is due to the downregulation of TE silencers. In fact, vertebrates have evolved a complex set of epigenetic modules to control TEs, such as KRAB-ZFPs, DNA methylation, small RNAs (piRNAs), and histone modifications [42]. We therefore systematically screened the expression patterns of these TE silencing systems during EHT. Surprisingly, the majority of TE silencers were downregulated in pre-HECs against VECs and HSCs (for example, 84.07% and 77.49% of KRAB-ZFPs in human and mouse, respectively) (Fig. 4C, F; Additional file 7: Table S9 and S10; KRAB-ZFP genes are available from [63]). Among the downregulated KRAB-ZPFs, ZNF84, ZNF382 and ZNF429 were found to bind significantly to L1 superfamily in human [64]. In embryonic stem cells, ZNF91 and ZNF93 can respectively repress SVAs and L1 in human [65], while Zfp932 regulates ERVK in mouse. In human, there is also evidence that ZNF268, ZNF300 and ZNF589 are related to hematopoietic differentiation [66]. It can be observed that some co-factors recruited by KRAB-ZFPs, such as TRIM28 (KAP1), CBX3 (HP1) and SETDB1, also exhibited relatively low expression levels in pre-HECs (Fig. 4C, F). TE silencers closely related to KRAB-ZFPs, such as DNMTs and NuRD complex also showed low expression in pre-HECs (Fig. 4E, H). In addition, although the overall expression levels were low, the HUSH complex (HUSHs), P-element induced Wimpy testis-related genes (PIWIs) and some other TE silencers were also expressed relatively lower in pre-HECs than in other cell types (Fig. 4C, F). Therefore, it can be inferred that various TE silencers were relaxed by specific mechanisms, leading to transient activation of TEs during pre-HEC specification (Fig. 4D, G). Interestingly, after the pre-HEC stage, those TE silencers were upregulated to re-suppress the TE activity. This also explains why some members of the DNMT complex (such as DNMT1 [67] and EZH2 [68]) and NuRD complex (such as HDAC1 and HDAC2 [69]) are required for HSC formation [70].

**Figure 4.**
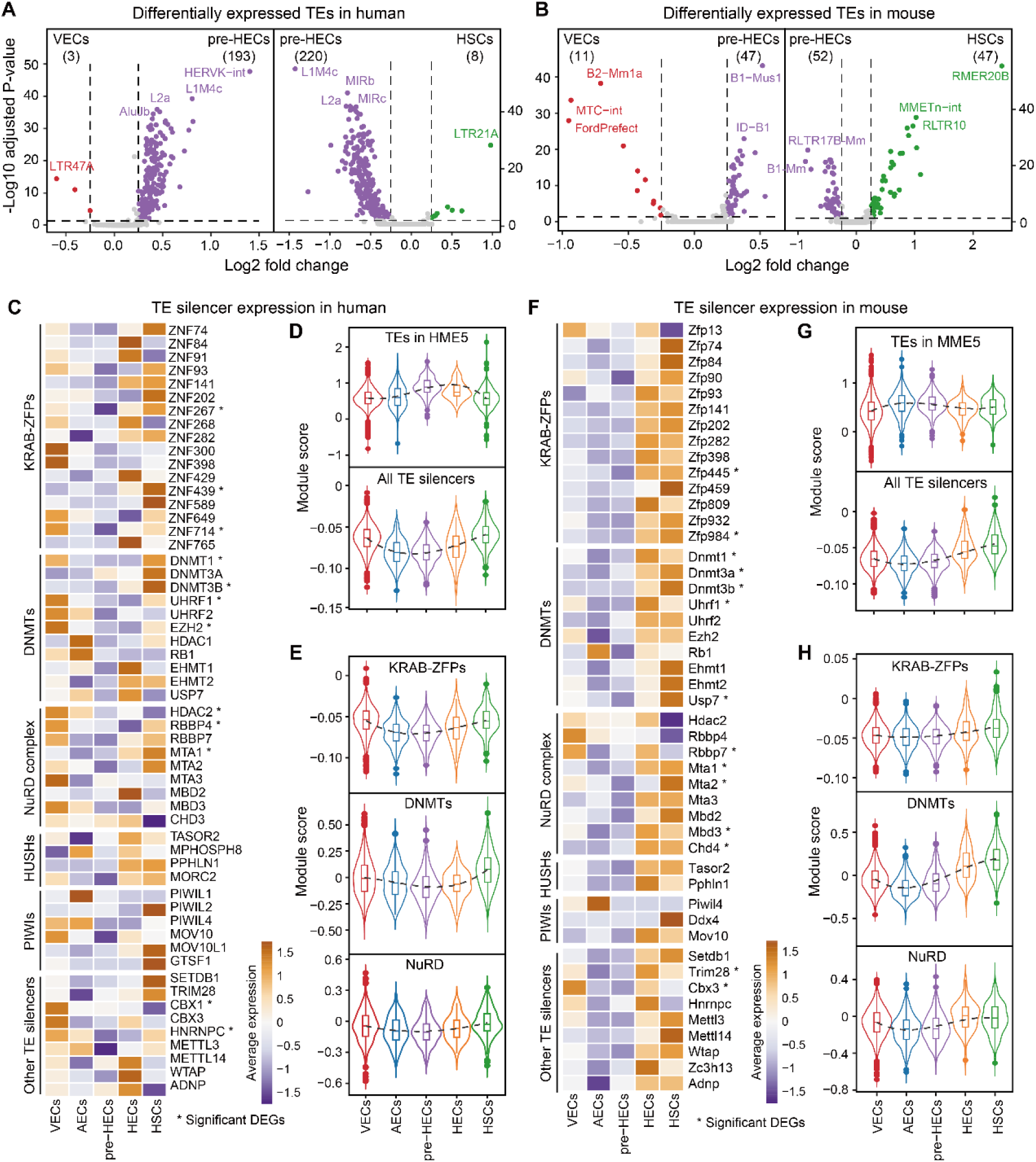
Expression of known TE silencers in human and mouse. **A, B** Differential expression of TEs in pre-HECs versus VECs and HSCs in human and mouse. **C, F** Expression heatmap of known TE silencing systems in human and mouse, including KRAB-ZFPs, DNMTs, NuRD complex, HUSHs, PIWIs and other TE silencers. **D, G** Expression trend of TEs (HME5 and MME5, kME ≥ 0.3) and all TE silencers. **E, H** Expression trend of specific TE silencing systems in human and mouse.

### TE product sensing facilitates immune activation during HSC orientation

It was found that massive TEs were activated at the pre-HEC stage, but the contribution of these TEs to the human and mouse EHT was unknown. Studies have shown that TEs are the main source of endogenous RNAs or cDNAs [39, 42]. The products transcribed from TEs are likely to activate downstream inflammatory and immune signaling pathways through pattern recognition receptors. We investigated genes that were more expressed in HECs/HSCs than in endothelial cells, and identified upregulation of a large number of RNA and DNA sensors both in human and mouse (Fig. 5A, E; Additional file 8: Table S1 and S2). IFIH1 (MDA5), DDX58 (RIG-I) of RLRs and NLRP1, NLRP2, NLRC3 and NLRX1 of NLRs seemed to be significantly upregulated in human HECs/HSCs. Although a few members of TLRs showed an upregulation trend in HECs/HSCs, the expression levels were low in both human and mouse (expression percentage is less than 10%). In addition, protein kinase R genes (PKRs) including EIF2A, EIF2AK1, EIF2AK2 and EIF2AK4 were both significantly upregulated in human and mouse HECs/HSCs. DNA sensors like the cGAS/STING (TMEM173) signals also appeared to be elevated in human and mouse HECs/HSCs, which were possibly activated by cDNA intermediates of retrotransposons (LINEs, SINEs and ERVs) [71, 72]. Typical downstream intermediates of RLRs and cGAS/STING, such as MAVS, TRAF3, TBK1, IRF3, and NF-κB, all showed highly conserved upregulation patterns in human and mouse HECs/HSCs (Fig. 5A, E; Additional file 8: Table S3 and S4). By functional enrichment analysis, both interferon alpha (IFNα) and interferon gamma (IFNγ) response pathways were detected to be enriched in human and mouse HECs/HSCs, confirming the activation of these PRRs (Additional file 8: Table S5 and S6). Interestingly, IFNAR1 and IFNGR2 consistently appeared to function earlier in human and mouse (immediately after TE expression upregulation), while IFNAR2 and IFNGR1 showed complementary patterns (Additional file 8: Table S3 and S4). Further gene set enrichment analysis (GSEA) on gene ontology (GO) showed that inflammatory signals (such as TNF and IL6) and immune response were enriched in human and mouse HECs/HSCs (Fig. 5B, F; Additional file 8: Table S5 and S6). Taking the above evidence together, we speculated that TE products (pervasively elevated in pre-HECs or earlier in partial AECs) could induce inflammatory signals through various PRRs during EHT, and trigger immune response pathways to activate HSC progression (Fig. 5C, G and D, H).

**Figure 5.**
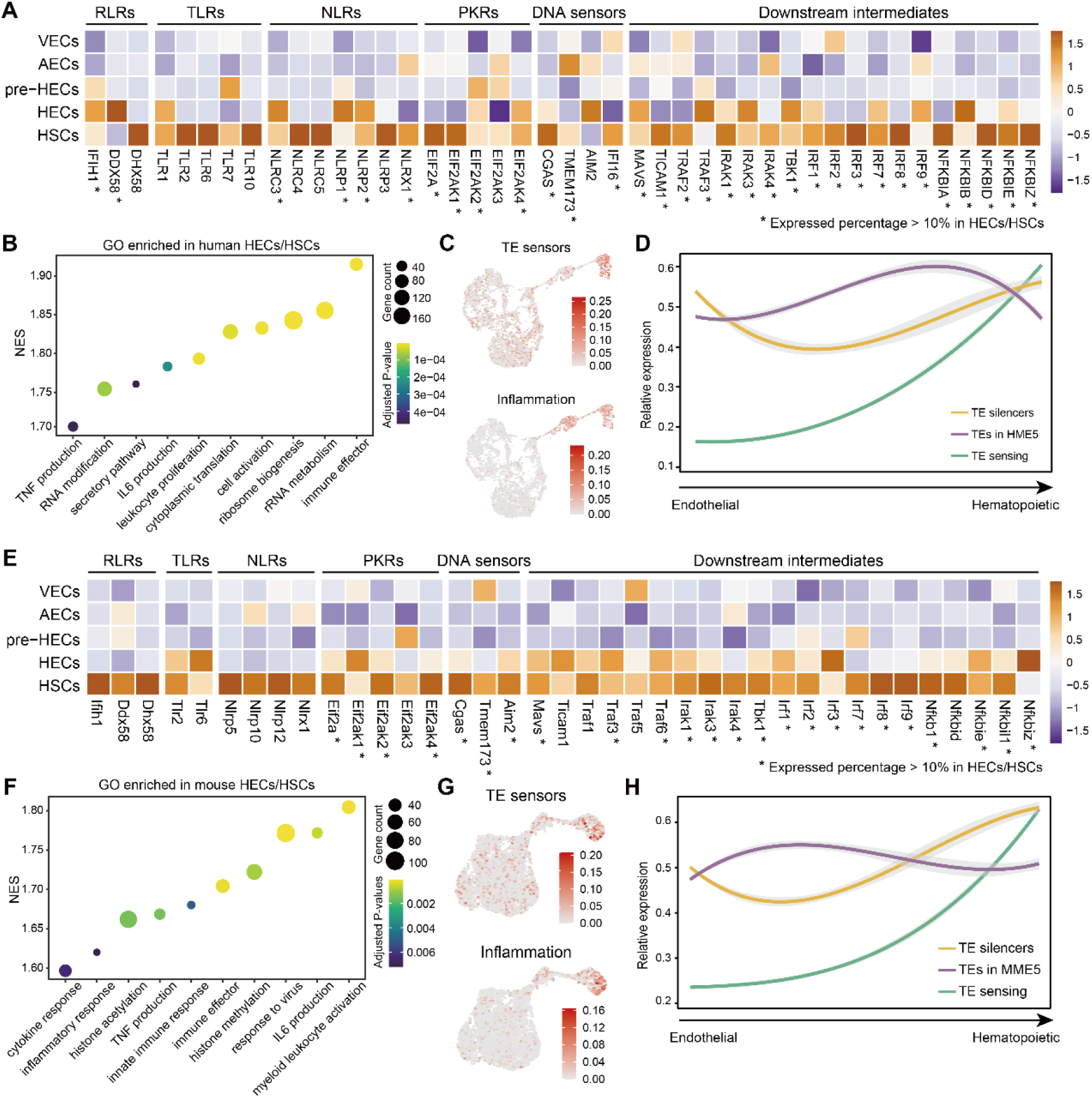
Expression of common TE sensors and functional enrichment for HECs/HSCs. **A, E** Expression heatmap of common TE sensors in human and mouse, including RLRs, TLRs, NLRs, PKRs, DNA sensors and downstream intermediates. **B, F** Gene set enrichment analysis of GO terms in human and mouse HECs/HSCs. **C, G** Module scores of TE sensors and inflammatory genes in human and mouse. **D, H** Expression trends of TE silencers, TEs (HME5 and MME5, kME≥0.3) and TE sensing genes during human and mouse EHT. The expression pattern of TEs is opposite to that of TE silencers, whereas TE sensors are less active until the HSC stage.

### TE accessibility is dynamically controlled during EHT

TEs are known to play important roles in development and disease processes as cis-regulatory elements [59]. To explore the potential cis-regulatory function of TEs on HSC origination, we systematically analyzed the scATAC-seq data in the E10.5 mouse AGM. Using cell types transferred from scRNA-seq data, a coherent EHT process can still be achieved based on scATAC-seq data (Fig. 6A; Additional file 9: Fig. S1A-D). The gene activities of specific EHT signatures obtained from scRNA-seq were well fitted to the developmental stage of EHT cell clusters (Fig. 6B; Additional file 9: Fig. S1E). For example, Gja5 showed pre-HEC-specific high activity on the UMAP embedding of scATAC-seq data. However, HECs were not able to be distinguished by scATAC-seq data, possibly due to the small number of captured cells. Next, we calculated the TE activities in each cell (reflecting the degree of TE accessibility) at the locus level. By applying differential accessibility analysis, we surprisingly found that TEs were more accessible in pre-HECs compared to endothelial cells (Fig. 6C; Additional file 10: Table S1), which aligned with our previous finding that TE expression is generally elevated in pre-HECs (Fig. 4A, B). While a total of 148 differentially accessible TEs (DATEs, average log2FC≥0. 25 & adjusted P-value≤0.05) were enriched in pre-HECs (Fig. 6C), only a few DATEs were identified between pre-HECs and HECs/HSCs (Additional file 10: Table S2). Notably, when differential accessibility analysis was performed on all peaks, we also found more differentially accessible peaks (DAPs) in pre-HECs compared to endothelial cells. This indicated that chromatin may undergo general regulation of accessibility, i.e., chromatin reprogramming, during the pre-HEC stage. In addition, TE accessibility did not change significantly when pre-HECs entered the HSC-committed stage, suggesting that the observed downregulation of TE expression in HECs/HSCs may not be influenced by chromatin accessibility. Unlike the low specificity of TE expression in scRNA-seq data, we identified a considerable number of cell type-specific open TEs in scATAC-seq data (Fig. 6D; Additional file 10: Table S3). Among them, AECs had the fewest specific open TEs (43), 16 of which overlapped with pre-HEC. DNAs accounted for the least amount of these cell type-specific open TEs, and LINEs accounted for more in pre-HECs than in other cell types (Fig. 6E). As expected, the majority of these open TEs played roles as distal enhancers.

**Figure 6.**
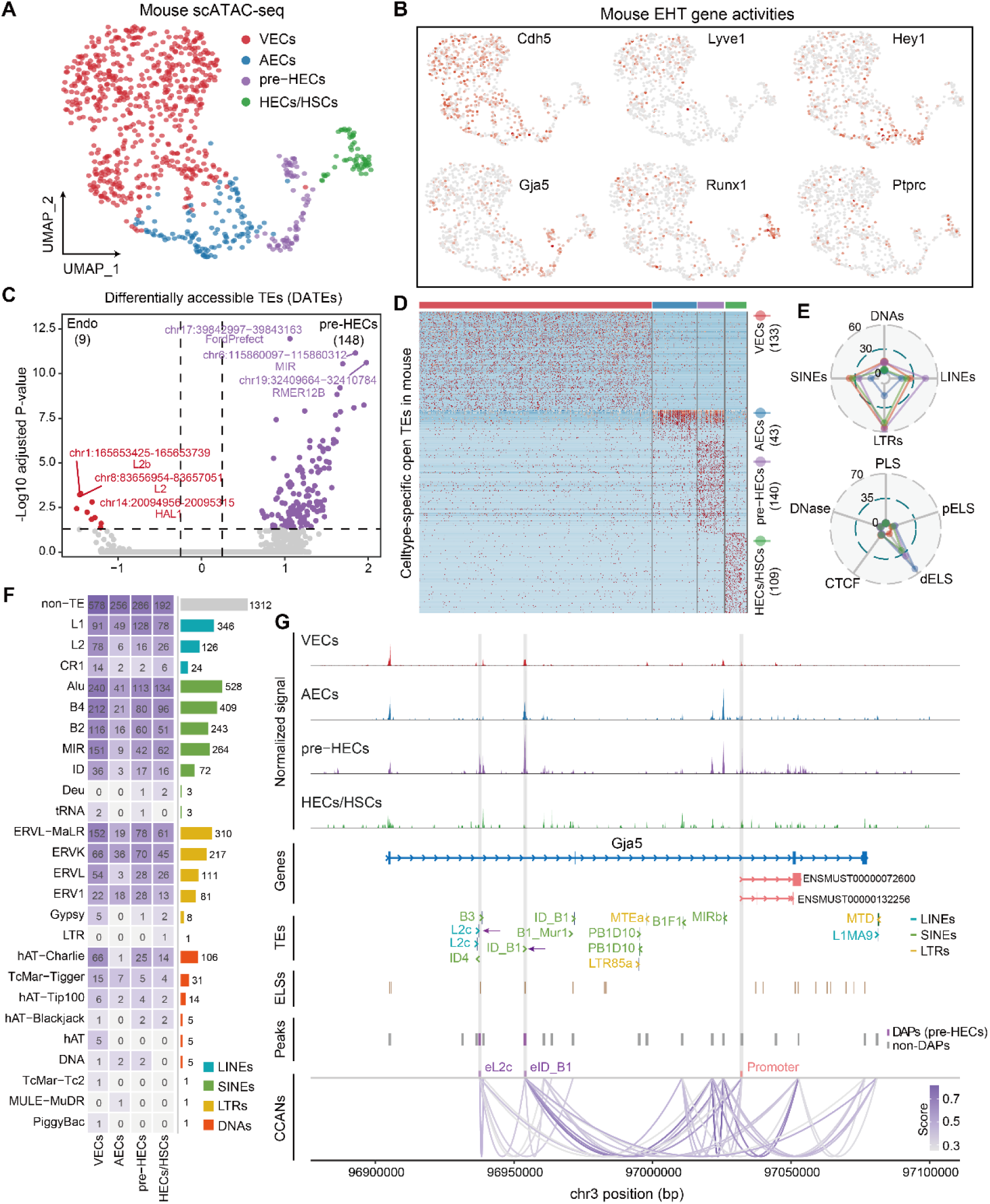
Mouse AGM scATAC-seq analysis and accessible TE identification. **A** Cell types recovered from mouse scATAC-seq data. **B** UMAP of EHT marker gene activities. Cdh5 marks endothelial cells, Lyve1 marks VECs, Hey1 marks AECs, Gja5 marks pre-HECs and partial AECs, Runx1 marks HECs/HSCs and Ptprc marks HSCs. **C** Differential accessible analysis of TEs in pre-HECs versus endothelial clusters (VECs and AECs). **D** Cell type-specific open TEs identified in mouse EHT. **E** TE composition and potential cis-regulatory functions of cell type-specific open TEs. **F** Overlaps of all cell type-specific accessible peaks with different TE superfamilies. **G** Tracks of normalized signals in each cell type, genes, TEs, ELSs, peaks and CCANs. Two potential enhancers derived by TEs and the promoter of Gja5 transcripts are highlighted with light grey bars.

Considering that TEs may function together with other accessible regions, we re-performed differential accessibility analysis for all peaks and identified a total of 4,230 cell type-specific DAPs, of which 2,918 overlapped with TEs, named TEPs (Fig. 6F; Additional file 10: Table S4). It can be seen that SINEs (especially Alu superfamily) exhibit relatively higher regulatory potential in each cell type. The gene regions closest to these TEPs contain many EHT-associated signatures, such as Gja4 (AECs), Gja5, Edn1 (pre-HECs), and Gata2, Cd44, Runx1 (HECs/HSCs) (Additional file 9: Fig. S2; Additional file 10: Table S4). Gja5 is a member of the connexin gene family, which had elevated expression in late AECs and pre-HECs (Fig. 2B, F). It can be noticed that the promoter of two transcripts (ENSMUST00000072600 and ENSMUST00000132256, annotated in EPD [73]) of Gja5 was specifically more accessible in pre-HECs (Fig. 6G), while two upstream promoter-like peaks showed high accessibility in both AECs and pre-HECs, which may permit the earlier expression of Gja5 observed in AECs. In addition, we found that two TEs (chr3:96937220−96937315-L2c and chr3:96954448−96954510: ID_B1) inside the gene body of Gja5 may function as enhancers (termed eL2c and eID_B1) to promote Gja5 expression in pre-HECs. The regions where these two TEs located are also annotated as ELSs in ENCODE. We applied Cicero [74] to predict the cis-co-accessibility networks (CCANs) among peaks detected near or inside Gja5. Although the potential enhancer eID_B1 had the greatest increase in accessibility in pre-HECs, it was also open in AECs and may interact with the two upstream promoter-like regions (Fig. 6G). Interestingly, the potential enhancer eL2c was only opened in pre-HECs, consistent with the accessibility pattern of the proximal promoter, and thus could be more likely to cooperatively increase the expression of Gja5. However, why Gja5 is upregulated in the pre-HEC stage of EHT remains to be further explored, although a recent study has pointed out its importance for HSCs to dampen oxidative stress [75].

### Cell type-specific accessible TEs shape hematopoietic cis-regulatory networks

It has been found that some TEs can act as enhancers to drive the expression of some hematopoietic-related genes, but the co-regulation mode of these TEs and the corresponding biological functions in the EHT process remain unclear. We used Cicero to construct the CCANs of all cell type-specific DAPs (including TEPs and non-TEPs) by filtering out links with co-access score less than 0.4 (Fig. 7A; Additional file 10: Table S5). Analysis of TE compositions of cell type-specific TEPs revealed that ID_B1 (Alu superfamily) was abundant in all cell types. The top-ranked TEs seemed to have high consistency across cell clusters, but they were enriched to different motifs in different EHT stages (Fig. 7B; Additional file 10: Table S6), which may be related to the variation accumulated on different copies of TEs during evolution [76, 77]. Surprisingly, TEs almost participate in shaping all cis-regulatory networks closely related to EHT process. For example, SOX and GATA binding sites were mostly open in VECs and RUNX binding sites gained increased accessibility in HECs/HSCs. We conducted a joint analysis of the enriched motifs and the corresponding transcription factors (TFs) (Fig. 7B, C), and found that although the motifs such as KLF (Klf7, Klf10 and Klf12) were active in AECs and later stages, the expression of these TFs were downregulated to control the developmental fate of AECs. In pre-HECs, the SOX motifs significantly increased the activity in AECs in advance, but the expression of TFs (Sox4, Sox6, Sox13 and Sox17) peaked after entering the pre-HEC stage. Similar to KLF, Gata3 and Gata6 had higher motif activities in both pre- HECs and HECs/HSCs, but were only highly expressed in pre-HECs. This dual regulation via motif binding activity and TF expression precisely shapes the lineage determination and functional specification during EHT.

**Figure 7.**
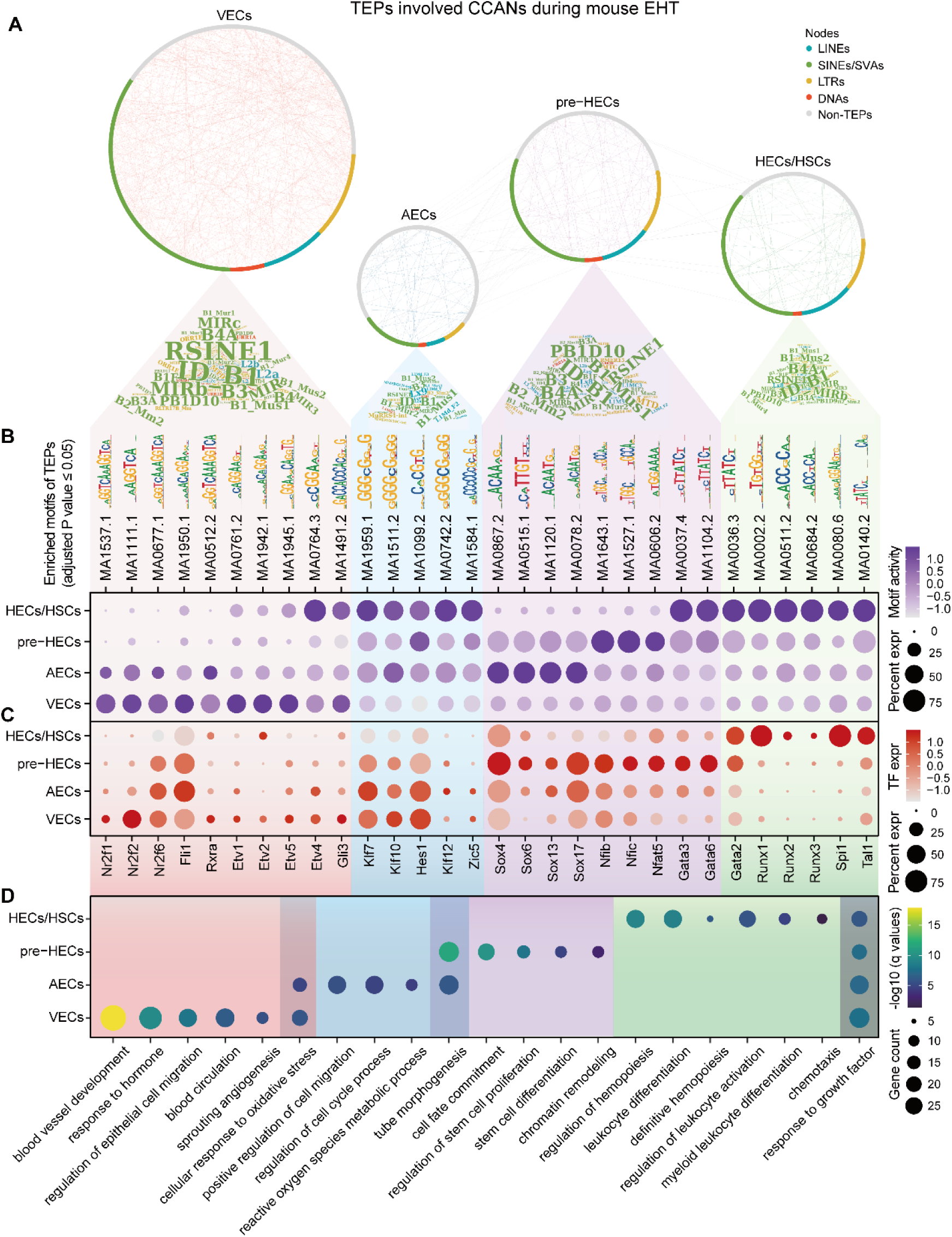
Cis-regulatory network analysis of TE-associated accessible peaks. **A** Cell type CCANs predicted by Cicero. TE families in each cell type are displayed as word clouds. **B** Enriched motifs of TE-associated accessible peaks in each cell type. **C** TF expression corresponding to motifs in (**B**). The expression data is extracted from scRNA-seq data. **D** GO enrichment of TF and target genes in each cell type. Overlaps are GO terms common to two or more cell types.

To explore the function of the TFs bound by TEPs, we predicted the cell type-specific TF-target network based on interactions from TRRUST [78]. Surprisingly, some EHT signatures were involved downstream of those TE-bound TFs (Additional file 9: Fig. S3; Additional file 10: Table S7), such as Kdr, Flt1 (VECs), Smad6, Vegfc (pre-HECs) and Kit, Ikzf1 (HECs/HSCs). The results of GO enrichment analysis also showed that these cis-regulatory networks shaped by cell type-specific TEs were enriched in various important functional modules during EHT (Fig. 7D; Additional file 10: Table S8), such as blood vessel development and tube morphogenesis in VECs and AECs, stem cell differentiation and chromatin reprogramming in pre-HECs, and regulation of hematopoiesis in HECs and HSCs.

### The hypoxic AGM niche may be partially responsible for the transient TE activation preceding hematopoietic fate commitment

The downregulation of TE silencing systems in pre-HECs may be the main reason for the enhanced TE activity, however the underlying mechanisms regulating these TE silencers remain unclear. Therefore, we focused on analyzing the specific gene expression and functional status of pre-HECs. The RNA velocity map showed that pre-HECs had a prominently high differentiation rate and are not in the cell cycle (Fig. 8A). In addition, more unspliced RNAs were found in pre-HECs (Fig. 8B), explaining the disordered direction of RNA velocity around pre-HECs (Fig. 2A). GSEA analysis showed that mRNA splicing, RNA catabolic process and cell cycle were downregulated in the pre-HEC stage (Fig. 8C; Additional file 11: Table S1). The downregulation of oxidative phosphorylation and upregulation of lipid metabolic process suggested that pre-HECs may undergo metabolic reprogramming [62]. Strikingly, genes related to epithelial to mesenchymal transition (EMT) and response to hypoxia were upregulated in pre-HECs. In cancers, hypoxia has been widely introduced to induce the EMT process and promote tumor cell metastasis [79, 80]. In zebrafish, hypoxia has been shown to strongly promote HSC formation through hypoxia-inducible factors (HIFs, hif-1a and hif-2a) and Notch signaling [81], which coincides with a study that induced human embryonic stem cells (hESCs) towards HSPC-like cells through a hypoxia differentiation system *in vitro* [82]. By staining the mouse embryo (E10) with a hypoxia indictor Pimonidazole (hypoxyprobe), it was also directly observed that the IAHC cluster region was hypoxic [83, 84]. We analyzed hypoxia-related genes on human EHT and found that EPAS1 (also known as HIF2A) and HIF3A [85] were highly expressed on pre-HECs, while HIF1A was more expressed on VECs and AECs (Fig. 8D; Additional file 11: Table S2). Many hypoxia-induced downstream genes were also found to be enriched in pre-HECs, such as SLC2A3 [86], CXCL12/CXCR4 [87], NOTCH1, VEGFC, EDN1, MMP2/MMP14, GATA6 TGFB2 and THBS1, etc. Spatial transcriptome analysis of human embryo (CS15) demonstrated that the expression of the above hypoxia-induced genes was also enriched in the adjacent region of IAHCs (Fig. 8F). Furthermore, we observed that the expression patterns of hypoxia-induced genes were exactly opposite to those of TE silencers (Fig. 8D, E), especially DNMT1 and UHRF1 (Fig. 8D, F). However, to the best of our knowledge, it has not been explored whether hypoxia can induce TE activation, although it is widely recognized that TEs play an important role in stress response [50, 88, 89]. Therefore, we first recalculated the TE expression landscape in the human AGM dataset. The results indicated that there seemed to be some other local hypoxic areas around AGM besides the pre-HECs, including the stromal cells, which also exhibited higher TE expression levels (Additional file 12: Fig. S1A-C). Few cell type-specific TEs were identified for each cell type in the AGM region (Additional file 12: Fig. SD), which is consistent with findings during the EHT trajectory (Fig. 2C, G). Besides, it was noticed that the expression pattern of HIF3A was closer to that of TE in various cell types in the AGM region (Additional file 12: Fig. S1B). HIF1A expressed in various cell types, whereas EPAS1 was enriched in endothelial cells (Additional file 12: Fig. S1E). Both stromal cells and epithelial cells had a certain degree of HIF3A expression, which could be part of the reason for their relatively higher TE expression levels (Additional file 12: Fig. S1A, B).

**Figure 8.**
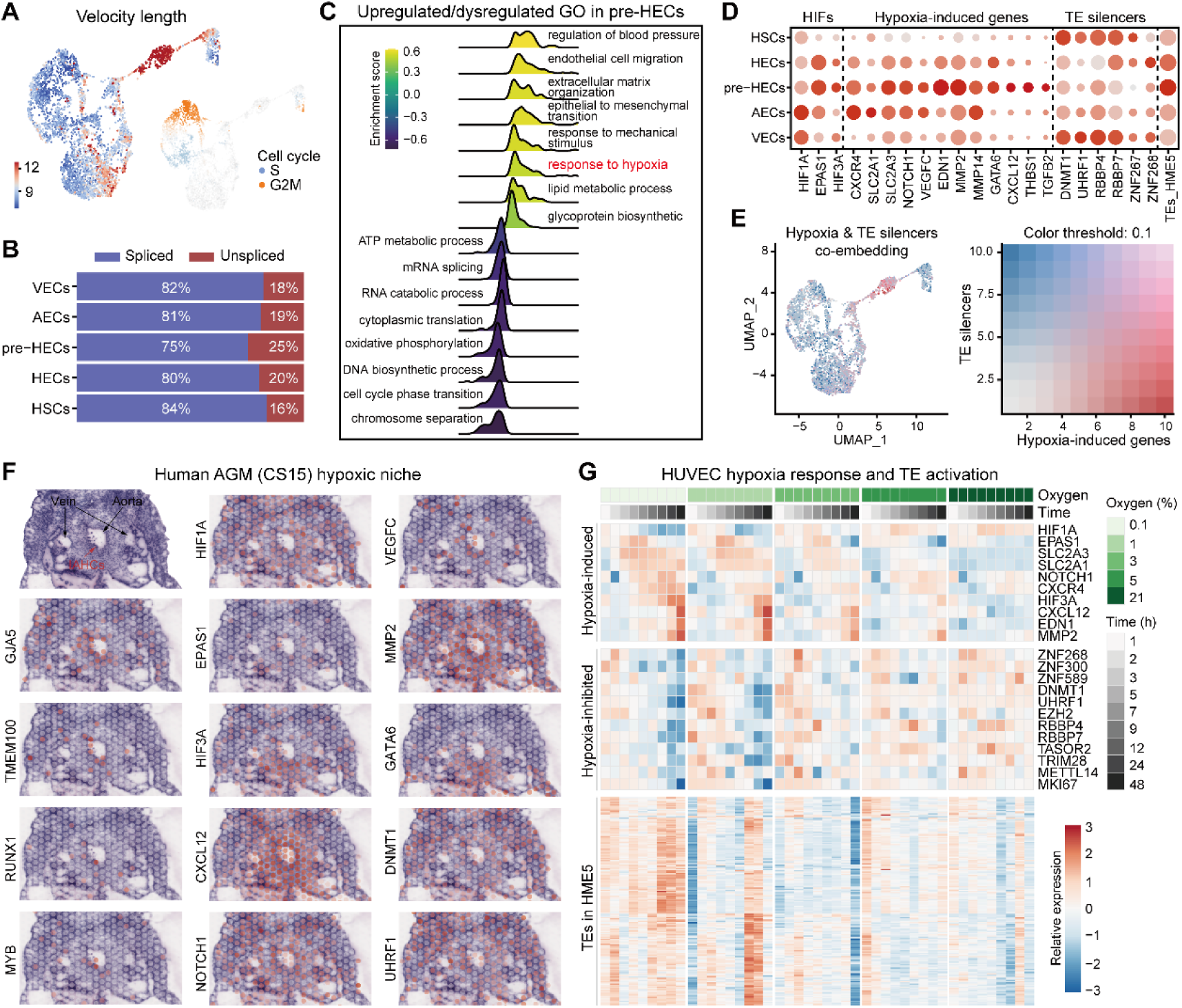
Hypoxia state analysis of pre-HECs and the AGM niche. **A** Velocity length and cell cycle scores on the human EHT UMAP. **B** The proportions of spliced and unspliced RNAs in each cell type. **C** Gene set enrichment analysis of GO terms in human pre-HECs **D** Expression of hypoxia-related genes and TE silencers in human EHT. **E** Co-embedding of expression of hypoxia-related genes and TE silencers. The expression patterns of the two seem to be opposite. **F** Spatial expression of hypoxia-related genes in the human AGM. **G** Expression heatmap of hypoxia-related genes, potential hypoxia-inhibited genes (TE silencers) and TEs in HME5 (kME≥0.3).

Furthermore, here we included a comprehensive time-series RNA-seq study [90] examining the hypoxia response of human umbilical vein endothelial cells (HUVECs) to confirm the relationship between hypoxia and TE activation. Excitingly, it is observed that TEs began to be broadly activated after 12 hours under extremely low oxygen concentrations (0.1% and 1%), whereas no significant upregulation of TEs was observed in groups with oxygen concentrations greater than 3% (Fig. 8G; Additional file 11: Table S3). Different TE classes showed similar upregulation patterns under hypoxia condition (Additional file 12: Fig. S2). Coincidentally, many TE silencers (such as KRAB-ZFP members ZNF268, ZNF300, ZNF589) were greatly downregulated after 12 hours of hypoxic culture. Interestingly, HIF3A still appears to be more correlated with TE expression patterns than HIF1A and EPAS1 (Fig. 8G). Collectively, we hypothesized that the hypoxic AGM niche might induce transient TE activation in pre-HECs by inhibiting the expression of TE silencers, which is postulated to be critical for the EHT process (Fig. 5E, J). By analyzing the expression of pre-HEC-specific markers in HUVEC data, it is observed that SOX17, HEY1 and HEY2 were not upregulated under hypoxia induction (Additional file 12: Fig. S2E, Fig. S3), suggesting that these genes may play distinct roles during pre-HEC specification.

## Discussion

TEs are abundant in the eukaryotic genomes and mounting evidence suggests that they have evolved essential roles in transcriptional and epigenetic regulation [42, 43, 57, 59, 76]. Recent single cell sequencing technologies to characterize the transcriptomes and epigenomes have revealed the broad expression and crucial roles of TEs in the developing embryos [47-49]. At present, although TEs have been revealed to be specifically expressed during definitive hematopoiesis and HSC regeneration [36, 50, 51], the underlying mechanisms of TE activation have not yet been elucidated. In addition, TEs are known to harbor a wide range of binding sites with regulatory potential (Fig. 1D, H), but their cis-regulatory roles during the EHT process remains to be investigated. In this work, we conducted a comprehensive analysis to understand the potential functionality of TEs during the definitive hematopoiesis in human and mouse at the single cell resolution. We demonstrated how cells conservatively program the EHT process and drive HSC formation by dynamically regulating the expression and chromatin accessibility of TEs. Finally, we deduced that the local hypoxic niche in AGM might be one of the important factors for the unique developmental state and TE activation in pre-HECs.

Leveraging the public single cell datasets of human and mouse AGM [22, 23], we reconstructed the EHT trajectories and presented the landscape of dynamic TE expression. Similar to the study in zebrafish [36], while only a few cell type-specific TEs (mainly LTRs) were identified during EHT (Fig. 2C, G), we unexpectedly observed that clusters of TEs were consistently upregulated during pre-HEC specification (Fig. 3B; Fig. 4A, B). The upregulation of TEs in mouse seems to occur already in the late stage of AECs (Fig. 3D). Coincidentally, TE silencing systems (including KRAB-ZFPs, DNMTs, NuRD complex, and HUSHs, etc.) [42, 44, 64, 66] were at relatively low levels from AECs to pre-HECs (Fig. 4C-H), which could at least partially account for the principle of TE activation in this period. Interestingly, two RNA transferases METTL3 and METTL14, which can form nuclear complex and control TE activity through m6A modification [91, 92], were also downregulated in pre-HECs.

Previous studies have shown that TE products can activate inflammatory signals through RLRs and promote HSC formation [36] and regeneration [50]. In contrast, we screened the common PRRs exhaustedly and found that except RLRs, many other PRRs were also upregulated in HECs/HSCs (Fig. 5A, E). For example, NLR family member X1 (NLRX1) [93] is conservatively enriched in both human and mouse HSCs, but its role in developmental hematopoiesis has not been described. DNA receptors (such as cGAS/STING) [72] also appeared to be activated after the pre-HEC stage, suggesting the possible presence of dsDNA from TEs or mitochondria. Many factors mediating these TE sensors and downstream inflammatory signaling were also specifically turned on in HECs/HSCs. It should be pointed out that dsRNAs from other sources, or some non-coding RNA, can also activate these PRRs [36, 39]. Notably, TE activation and TE sensing are not synchronized, that is, TEs that are largely transcribed in pre-HECs do not activate PRRs and inflammatory signals until HECs/HSCs. A possible explanation for this might be that the RNA catabolic process and cell metabolism are quiescent in pre-HECs (Fig. 8B, C), which delays the TE sensing (Fig. 9). Such TE activation and sensing is conserved in human and mouse, which resembles a rehearsal mechanism during the EHT process. After hematopoietic differentiation, nascent HSCs learn antigen-like properties from endogenous nucleic acid repertoire (e.g. TEs) and complete a pluripotent immune activation, which lays the foundation for the anti-pathogen ability of various immune cells differentiated from mature HSC.

**Figure 9.**
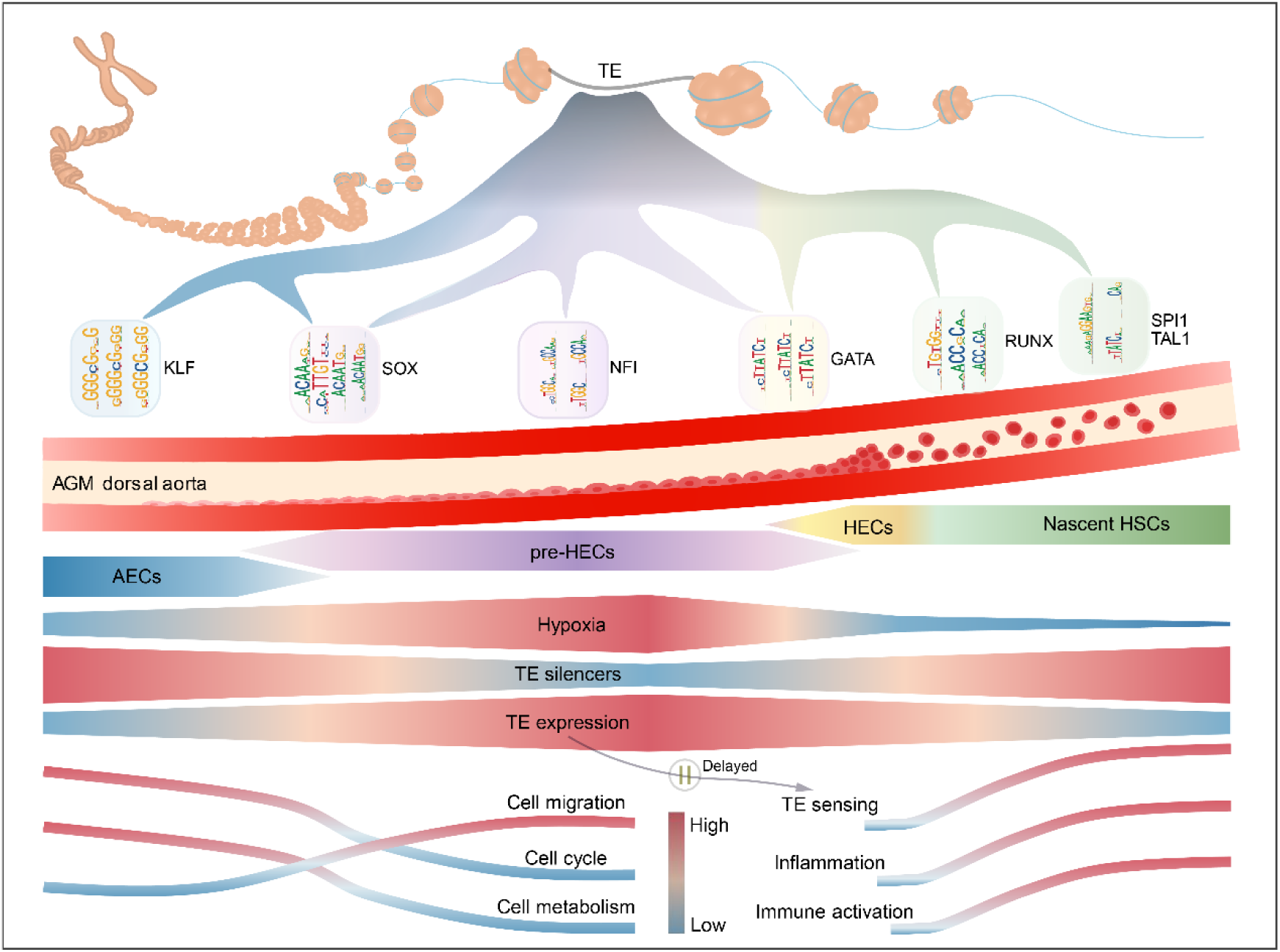
Schematic diagram of dynamic regulations of TEs during EHT. The EHT process in the AGM dorsal aorta is shown in the middle vessel. The process of TEs driving EHT by providing different cis-regulatory elements through dynamic accessibility is presented above. Below the blood vessel, the AGM hypoxic niche induces TE activation in pre-HECs and triggers delayed TE sensing and inflammatory signaling through pattern recognition receptors, thereby promoting the formation of nascent HSCs.

Cis-regulatory elements, especially enhancers, are another important way in which TEs exert genome regulatory functions [57, 59, 61, 76, 77]. In this study, we observed that TEs show cell type-specific dynamic accessible patterns (Fig. 6D). The distribution of TE-related open regions was detected near many EHT marker genes (Additional file 9: Fig. S2). We identified two TE-associated enhancers, eL2c and eID_B1 (upstream of the Gja5 transcript), whose accessibilities were specifically increased in pre-HECs and were predicted to interact with the Gja5 promoter (Fig. 6G). However, the transcription factors that bind them have not been determined. Through motif prediction and TF expression analysis of cell type-specific TEPs, we recovered the complete TE-involved cis-regulatory networks during EHT (Fig. 7). Interestingly, we found that the activity of many motifs and the expression of corresponding TFs are not fully synchronized. The open range of motifs seems to be wider, while the TF expression is more specific. For example, some key TF binding motifs (such as SOX, GATA) appear to open earlier, while the TF expression is tightly timed (Fig. 7B, C; Fig. 9).

What are the conditions for the activation of TEs during pre-HEC specification, and whether the AGM niche has an effect on the TE activation? These are the objects of much speculation. Many evidences show that environmental stress (including heat shock, oxidative and chemotherapy) [50, 94] is one of the driving forces to induce TE activation. In our study, we inferred that the AGM hypoxic niche may be at least partially responsible for the activation of TEs in pre-HECs. Especially after the elevated expression of hypoxia-inducible factors EPAS1 and HIF3A, the TE silencers were downregulated, accompanied by the increase of TE expression (Fig. 8D, E). Indeed, hypoxia in the AGM region (IAHC cluster) has already been observed and shown to promote HSC formation [81-84]. Our study supports these findings and attributes the role of hypoxia on HSC development in part to TE activation. Interestingly, the stromal cells and epithelial cells in AGM were also hypoxic to some extent, and TE expression was also relatively high (Fig. 8F; Additional file 12: Fig. S1A), indicating that the activation of TEs by hypoxia appears to be cell type insensitive. We recognize that HIF3A seems to be more related to the TE expression, as evidenced by the upregulation of TEs in stromal and epithelial cells that do not express EPAS1 but express HIF3A (Additional file 12: Fig. S1B, E). Furthermore, by analyzing a time-series RNA-seq study examining the hypoxia response of HUVECs, we observed that TE silencers were downregulated and the TE expression was broadly upregulated after 12 h of hypoxic culture, further confirming our hypothesis (Fig. 8G). In fact, the effects of hypoxia on HSC development are multifaceted, for example, we found that cell cycle and metabolism were inhibited in hypoxic pre-HECs (Fig. 8C; Fig. 9). Conversely, genes related to cell migration were upregulated, which matches the findings in tumor EMT [95, 96].

## Conclusions

TEs are domesticated during the evolution of eukaryotic genomes and mediate the emergence of novel regulatory elements [42]. TEs have been found to be specifically expressed to promote definitive hematopoiesis and HSC regeneration through RLRs and inflammatory signaling [31, 36, 50, 51]. Our study not only extends the potential upstream and downstream of TE transcription during EHT at the single-cell level, but also fills in the gap of TEs as cis-regulatory elements driving HSC development. We found that TEs were massively upregulated during pre-HEC specification, coinciding with the downregulation of TE silencers at this stage. PRRs-mediated TE product sensing and activation of inflammatory signaling are delayed until the HSC stage. These observations are highly conserved between human and mouse. Analysis of scATAC-seq data reveals that dynamically accessible TEs shape the hematopoietic cis-regulatory network to coordinate the EHT process. We additionally reported that the hypoxic AGM niche may be partially responsible for the transient TE activation before hematopoietic fate commitment. Further investigations are required to confirm such a hypothesis. In summary, this study provides a systematical single-cell analysis to uncover how TEs, through dynamic expression and chromatin accessibility, orchestrate the EHT process and drive HSC formation.

## Methods

### TE coverage, distribution, and regulatory potential analysis

The TE annotation data of human (hg38) and mouse (mm10) were obtained from the UCSC Genome Browser database (https://genome.ucsc.edu/) [97]. The genomic annotations (intergenic, intron, 3’ UTR, 5’ UTR, CDS) and unmasked CpG islands data were also downloaded from UCSC. The intersection of TEs and gene structures was measured using BEDTools (v2.30.0) [98]. ChIPseeker (v1.34.1) [99] was used to visualize the TE distributions with respect to protein-coding genes. The cCREs annotations (including PLS, pELS, dELS, CTCF-only and DNase-H3K4me3) were downloaded from ENCODE [54]. To improve the annotation accuracy of TE regulatory potential, the overlapping of TE and cCRE was required to be more than 50% of the TE length. The heatmap representations were generated using the R package pheatmap (v1.0.12).

### Single cell RNA-seq data processing

The single-cell raw sequencing data of human and mouse AGM were downloaded from GEO (https://www.ncbi.nlm.nih.gov/geo/) with accession numbers GSE162950 [23] and GSE137117 [22]. The detailed information of samples used in this study can be found in Additional file 3: Table S1. Reads were mapped to the human (refdata-gex-GRCh38-2020-A) and mouse (refdata-gex-mm10-2020-A) reference genomes using CellRanger (v7.1.0). The R package Seurat (v4.3.0) [100] was used to perform downstream analysis. Cells with less than 200 unique molecular identifiers (UMIs) or greater than 15% mitochondrial expression were removed and clusters with unusual low RNA features or counts were also filtered in further analysis. Batch effects were corrected by Harmony (v0.1.1) [101]. SCTransform (v0.3.5) [102] was used to normalize the clean data followed by dimension reduction and clustering through Seurat. Marker genes were identified using FindAllMarkers with MAST [103]. Cell types were annotated according to the marker genes provided in [23] (Additional file 4). Integration of the human and mouse EHT data were achieved by Seurat CCA based on the shared homologous genes. The R package biomaRt (v2.54.1) [104] was used to map gene symbols of mouse to human.

### EHT trajectory reconstruction

To reconstruct the EHT trajectory, Velocyto (v0.17.17) [105] was applied to estimate the RNA velocity from the bam files generated by Cell Ranger. The velocity maps were visualized by scVelo (v0.2.5) [106]. The latent time was estimated using dynamical modeling that models the full splicing kinetics in scVelo. Cell cycle scores along EHT trajectory were calculated based on phase marker genes [107].

### Single cell TE quantification and differential expression analysis

We applied scTE (v1.0) [49] to quantify the TE expression at the family level in human and mouse scRNA-seq data. To keep the consistency of the read counting results, we incorporated the same gene annotations as Cell Ranger and TE annotations from UCSC to build the genome indices. Count matrix of only LINEs, SINEs/SVAs, LTRs and DNAs were kept and merged into the Seurat object for further analysis. Cell type-specific marker TEs were identified using FindAllMarkers with default parameters. Differential expression analysis of TEs was performed using FindMarkers with default parameters. The specificity of a TE was measured by the percent difference in expression of the TE between the target cell type and other cell types.

### Co-expression gene and TE module analysis

Cell type-specific marker genes (average log2FC≥0.5 and adjusted P-value≤0.05) and TEs counting more than 50 were extracted for weighted gene co-expression network analysis (WGCNA) using hdWGCNA (v0.2.16) [108], which extends the standard WGCNA [109] pipeline into scRNA-seq analysis. The single cells were first aggregated into pseudobulk (meta) cells to reduce the drop out effect. The co-expression networks were visualized with UMAP. The module connectivity score (kME) was computed based on module eigengene. The module scores of TEs with kME≥0.3 in HME5 and MME5 were calculated using AddModuleScore in Seurat.

### TE silencing and sensing analysis

Genes related to TE silencing were collected from the literature [42, 44]. Potential KRAB-ZFP genes in human and mouse were obtained from [63]. The whole list of TE silencers analyzed in this study can be found in Additional file 7: Table S9 and S10. TE silencers with lower mean expression in pre-HECs than in other cell types were selected and displayed as heatmaps. Genes associated with TE sensing (including PRRs and downstream intermediates) were also extracted from publications [35, 37, 39, 71, 72]. TE sensing genes and inflammatory factors are listed in Additional file 8: Table S3 and S4. Differential expression analysis of genes between hematopoietic cells (HECs/HSCs) and endothelial cells (VECs/AECs) were performed using FindMarkers with default parameters. The module scores of different gene sets were calculated using AddModuleScore.

### Functional enrichment analysis

The gene set enrichment analysis (GSEA) was performed using clusterProfiler (v4.6.2) [110]. Both Gene Ontology (biological process) and Molecular Signatures Database (hallmark gene sets) are included. Genes were ranked according to fold changes calculated in Seurat.

### Single cell ATAC-seq data processing

The raw sequencing data of mouse AGM (E10.5) was downloaded from GEO with accession GSE137115 [22]. Reads were mapped to the mouse reference genome (refdata-cellranger-arc-mm10-2020-A) using cellranger-atac (v2.1.0). The R package Signac (v1.9.0) [111] was used to perform downstream analysis, including quality control, normalization, dimension reduction and clustering. After estimating the gene activities, the cell types of scATAC-seq data were annotated through cross-modality integration and label transfer from scRNA-seq data using CCA [102]. The final cell types were corrected according to the gene activities of known EHT markers.

### Single cell TE accessibility estimation and differential accessible analysis

The count matrix of TEs was estimated using FeatureMatrix in Signac. Cell type-specific open TEs were identified by FindAllMarkers. Differentially accessible peaks and TEs between cell types were identified using FindMarkers. Each of the open TEs was assigned to the closest gene using ClosestFeature. TE-related differentially accessible peaks (Additional file 9: Fig. S2) were plotted on the mouse genome using karyoloteR (v1.24.0) [112].

### Cis-co-accessible network (CCAN) construction

The cis-co-accessible peaks were identified using Cicero (v1.3.9) [74]. The links with coaccess score more than 0.4 were extracted to construct the CCANs. The CCAN network of all differentially accessible peaks was visualized in Cytoscape (v3.9.0) [113].

### Motif enrichment and TF expression analysis

The motif enrichment analysis was performed in Signac. The motif position frequency matrices were from JASPAR [114]. Motifs enriched in TE-related differentially accessible peaks were found by FindMotifs. The motif activity was computed by chromVAR (v1.20.2) [115]. Active motifs were selected by combining with the expression of corresponding TFs from scRNA-seq data. The cell type-specific TF-target network (Additional file 9: Fig. S3) was constructed based on interactions from TRRUST [78]. The average expression of the target genes in the target cell type were required to be more than 0.25.

### Spatial transcriptome data processing

The raw spatial transcriptome sequencing data of human AGM (CS15, sample 7) was downloaded from GEO with accession GSE162950 [23]. Reads were mapped to the human reference genome (refdata-gex-GRCh38-2020-A) using Space Ranger (v2.0.1). The R package Seurat was used to perform downstream analysis. The expression data were normalized using SCTransform.

### HUVEC bulk RNA-seq data processing

The raw sequencing data of HUVECs against hypoxia stress was downloaded from SRA (https://www.ncbi.nlm.nih.gov/Traces/study/) with accession PRJNA561635 [90]. The detailed information of samples used in this study can be found in Additional file 3: Table S2. We treated each sample as a single cell and thus can still use scTE to quantify TE and gene expression. Only wild type samples were included for further analysis. The gene modules of pre-HEC markers (Additional file 12: Fig. S3) in HUVEC data were predicted using WGCNA [109].

## Abbreviations

EHT: endothelial-to-hematopoietic transition;
TEs: transposable elements;
HSCs: hematopoietic stem cells;
ECs: endothelial cells;
AGM: aorta-gonad-mesonephro;
AECs: arterial endothelial cells;
HECs: hematopoietic endothelial cells;
IAHCs: intra-aortic hematopoietic clusters;
scRNA-seq: single-cell RNA sequencing;
scATAC-seq: single-cell sequencing assay for transposase-accessible chromatin;
TNF: tumor necrosis factor;
IFN: interferon;
PRRs: pattern recognition receptors;
TLRs: Toll-like receptors;
RLRs: RIG-I-like receptors;
NLRs: NOD-like receptors;
CLRs: C-type lectin receptors;
TRAFs: TNF receptor-associated factors;
ssRNAs: single-stranded RNAs;
dsRNAs: double-stranded RNAs;
DNAs: DNA transposons;
LINEs: long interspersed nuclear elements;
SINEs: short interspersed nuclear elements;
SVA: SINE-VNTR-Alu;
LTRs: long terminal repeats;
ERVs: endogenous retroviruses;
KRAB-ZFP: Krüppel-associated box zinc finger protein;
DNMT: DNA methyltransferases;
NuRD: nucleosome remodeling deacetylase;
HUSH: human silencing hub;
TSS: transcription start sites;
TTS: transcription termination sites;
cCREs: candidate cis-regulatory elements;
PLS: promoter-like sites;
pELS: proximal enhancer-like signatures;
dELS: distal enhancer-like signatures;
kME: module connectivity;
PIWIs: P-element induced Wimpy testis-related genes;
PKRs: protein kinase R genes;
GSEA: gene set enrichment analysis;
GO: gene ontology;
DATEs: differentially accessible TEs;
DAPs: differentially accessible peaks;
TEPs: TE overlapped DAPs;
CCANs: cis-co-accessibility networks;
EMT: epithelial to mesenchymal transition;
HIFs: hypoxia-inducible factors;
hESCs: human embryonic stem cells;
HUVECs: human umbilical vein endothelial cells;
UMIs: unique molecular identifiers;
WGCNA: weighted gene co-expression network analysis.

## Declarations

### Ethics approval and consent to participate

Not applicable.

### Consent for publication

Not applicable.

### Availability of data and materials

The scRNA-seq and spatial transcriptome data for human AGM that were analyzed in this study are available from GEO (GSE162950) [23]. The scRNA-seq and scATAC-seq data for mouse AGM are available from GEO (GSE137117) [22]. The time series RNA-seq data for HUVEC are available from SRA (PRJNA561635) [90]. The detailed information of samples used in this study can be found in Additional file 3. All analysis pipelines, in-house scripts and files for reproducing the results in this study can be accessed at https://github.com/ventson/hscTE. We also provide a web interface (https://bis.zju.edu.cn/hscTE, implemented using UCSC Cell Browser [116]) to visualize TE and gene expression during human and mouse EHT. The multi-faceted display (including TEs, CpG, cCREs, peaks and genome coverages) of mouse EHT scATAC-seq data is available from https://bis.zju.edu.cn/hscTE/jbrowse/?data=mouse, which is implemented by JBrowse [117].

### Competing interests

The authors declare that they have no competing interests.

### Funding

This work was supported by the National Key Research and Development Program of China (No. 2018YFC0310602; 2016YFA0501704), National Natural Sciences Foundation of China (No. 31771477; 32070677), the 151 Talent Project of Zhejiang Province (first level), the Fundamental Research Funds for the Central Universities, Jiangsu Collaborative Innovation Center for Modern Crop Production and Collaborative Innovation Center for Modern Crop Production co-sponsored by province and ministry, Doctor Foundation of The Second Hospital of Shanxi Medical University (No. 201701-4), Natural Science Foundation of Shanxi Province (No. 201901D111381).

### Authors’ contributions

MC, HH, and WDL conceived and supervised the study. CF, RXT, and SGX processed the data and prepared the manuscript. YHC helped RNA velocity and trajectory analysis. SDL and XTH helped the spatial transcriptome analysis. YCZ, YJL, YMH, YSH, HP, and ZXW contributed to the analysis pipeline construction and modification. HYC, SLZ, and QYN helped the scATAC-data analysis. JYH provided constructive discussion and suggestions. CF, RXT, SGX, MC, HH, and WDL wrote and revised the manuscript with input from all the other authors. All authors read and approved the final manuscript.

## Acknowledgements

The authors would like to thank members in Ming Chen’s lab for discussion and valuable suggestions.

## Authors’ information

Department of Bioinformatics, Zhejiang University College of Life Sciences, Hangzhou 310058, China

Cong Feng, Saige Xin, Yuhao Chen, Sida Li, Xiaotian Hu, Yincong Zhou, Yueming Hu, Yanshi Hu, Zexu Wu, Haoyu Chao, Shilong Zhang, Qingyang Ni, Ming Chen Bioinformatics Center, The First Affiliated Hospital, Zhejiang University School of Medicine, Hangzhou 310058, China

Cong Feng, Yongjing Liu, Jinyan Huang, Ming Chen

Bone Marrow Transplantation Center, the First Affiliated Hospital, Zhejiang University School of Medicine, Hangzhou 310058, China

Ruxiu Tie, He Huang

Liangzhu Laboratory, Zhejiang University Medical Center, Hangzhou 310058, China Ruxiu Tie, He Huang

Institute of Hematology, Zhejiang University, Hangzhou 310058, China Ruxiu Tie, He Huang

Zhejiang Province Engineering Laboratory for Stem Cell and Immunity Therapy, Hangzhou 310058, China

Ruxiu Tie, He Huang

Department of Hematology, The Second Clinical Medical College of Shanxi Medical University, Shanxi Medical University, Taiyuan 030000, China

Ruxiu Tie

Department of Hematology-Oncology, Taizhou Hospital of Zhejiang Province, Linhai 317000, China

Ruxiu Tie, Wenda Luo

Department of Veterinary Medicine, Zhejiang University College of Animal Sciences, Hangzhou 310058, China

Hang Pan

## Supplementary information

**Additional file 1: Table S1.** TE families, superfamilies and classes in human. **Table S2.** TE families, superfamilies and classes in mouse. **Table S3.** Genome coverage of TEs in human. **Table S4.** Genome coverage of TEs in mouse. **Table S5.** Genomic distribution of TEs in human. **Table S6.** Genomic distribution of TEs in mouse.

**Additional file 2: Table S1.** Overlaps among TEs and CpG islands in human. **Table S2.** Overlaps among TEs and CpG islands in mouse. **Table S3.** Overlaps among TEs and promoter-like sites (PLS) in human. **Table S4.** Overlaps among TEs and proximal enhancer-like sites (pELS) in human. **Table S5.** Overlaps among TEs and distal enhancer-like sites (dELS) in human. **Table S6.** Overlaps among TEs and CTCF-only sites in human. **Table S7.** Overlaps among TEs and DNase-H3K4me3 sites in human. **Table S8.** Overlaps among TEs and promoter-like sites (PLS) in mouse. **Table S9.** Overlaps among TEs and proximal enhancer-like sites (pELS) in mouse. **Table S10.** Overlaps among TEs and distal enhancer-like sites (dELS) in mouse. **Table S11.** Overlaps among TEs and CTCF-only sites in mouse. **Table S12.** Overlaps among TEs and DNase-H3K4me3 sites in mouse. **Table S13.** Copy number of cCREs overlapped with TEs in human. **Table S14.** Copy number of cCREs overlapped with TEs in mouse.

**Additional file 3: Table S1.** Human and mouse AGM datasets included in this study. **Table S2.** HUVEC bulk RNA-seq dataset (PRJNA561635) included in this study.

**Additional file 4: Figure S1.** Steps to reconstruct the human EHT trajectory. **Figure S2.** Steps to reconstruct the mouse EHT trajectory. **Figure S3.** Integration of the human and mouse EHT scRNA-seq data.

**Additional file 5: Table S1.** Marker TEs in human EHT. **Table S2.** Marker TEs in mouse EHT. **Table S3.** Marker genes in human EHT. **Table S4.** Marker genes in mouse EHT.

**Additional file 6: Table S1.** Co-expressed gene and TE modules in human. **Table S2.** Co-expressed gene and TE modules in mouse. **Table S3.** Common TEs in HME5 and MME5.

**Additional file 7: Table S1.** Differentially expressed TEs between pre-HECs and VECs in human. **Table S2.** Differentially expressed genes between pre-HECs and VECs in human. **Table S3.** Differentially expressed TEs between pre-HECs and HSCs in human. **Table S4.** Differentially expressed genes between pre-HECs and HSCs in human. **Table S5.** Differentially expressed TEs between pre-HECs and VECs in mouse. **Table S6.** Differentially expressed genes between pre-HECs and VECs in mouse. **Table S7.** Differentially expressed TEs between pre-HECs and HSCs in mouse. **Table S8.** Differentially expressed genes between pre-HECs and HSCs in mouse. **Table S9.** Average expression of TE silencers in human. **Table S10.** Average expression of TE silencers in mouse.

**Additional file 8: Table S1.** Differentially expressed genes in human hematopoietic cells (HECs/HSCs) vs endothelial cells (VECs/AECs). **Table S2.** Differentially expressed genes in mouse hematopoietic cells (HECs/HSCs) vs endothelial cells (VECs/AECs). **Table S3.** Average expression of pattern recognition receptors and downstream signals in human. **Table S4.** Average expression of pattern recognition receptors and downstream signals in mouse. **Table S5.** GSEA analysis of genes in human hematopoietic cells (HECs/HSCs) vs endothelial cells (VECs/AECs). **Table S6.** GSEA analysis of genes in mouse hematopoietic cells (HECs/HSCs) vs endothelial cells (VECs/AECs).

**Additional file 9: Figure S1.** Steps to reconstruct the mouse EHT trajectory from scATAC-seq data. **Figure S2.** Genome landscape of differentially accessible peaks (DAPs). **Figure S3.** The TF-target network in mouse EHT.

**Additional file 10: Table S1.** Differentially accessible TEs between pre-HECs and VECs/AECs. **Table S2.** Differentially accessible TEs between pre-HECs and HECs/HSCs. **Table S3.** Cell type-specific open TEs. **Table S4.** Closest genes and annotations of differentially accessible peaks in mouse EHT. **Table S5.** Co-access scores of cell type-specific peak pairs. **Table S6.** Enriched motifs of TE associated cell type-specific peaks (TEPs). **Table S7.** Activated TF-target pairs during EHT (TFs are corresponding to TE-related motifs). **Table S8.** GO enrichment results of cell type-specific network modules of TE-bound TFs and downstream targets.

**Additional file 11: Table S1.** GSEA analysis of pre-HECs vs other cell types. **Table S2.** Average expression of hypoxia-related genes in human EHT cell types. **Table S3.** Normalized counts of genes and TEs in the HUVEC dataset (PRJNA561635).

**Additional file 12: Figure S1.** The hypoxic niche and TE expression in human AGM scRNA-seq data. **Figure S2.** Expression heatmap of each TE family (grouped into four TE classes) in the HUVEC bulk RNA-seq data. **Figure S3.** Five gene modules of pre-HEC markers on the HUVEC data.

## Notes

https://bis.zju.edu.cn/hscTE

https://bis.zju.edu.cn/hscTE/jbrowse/?data=mouse

### Competing Interest Statement

The authors have declared no competing interest.

