## Additional file 4 for "Systematic single-cell analysis reveals dynamic control of transposable element activity orchestrating the endothelial-to-hematopoietic transition"

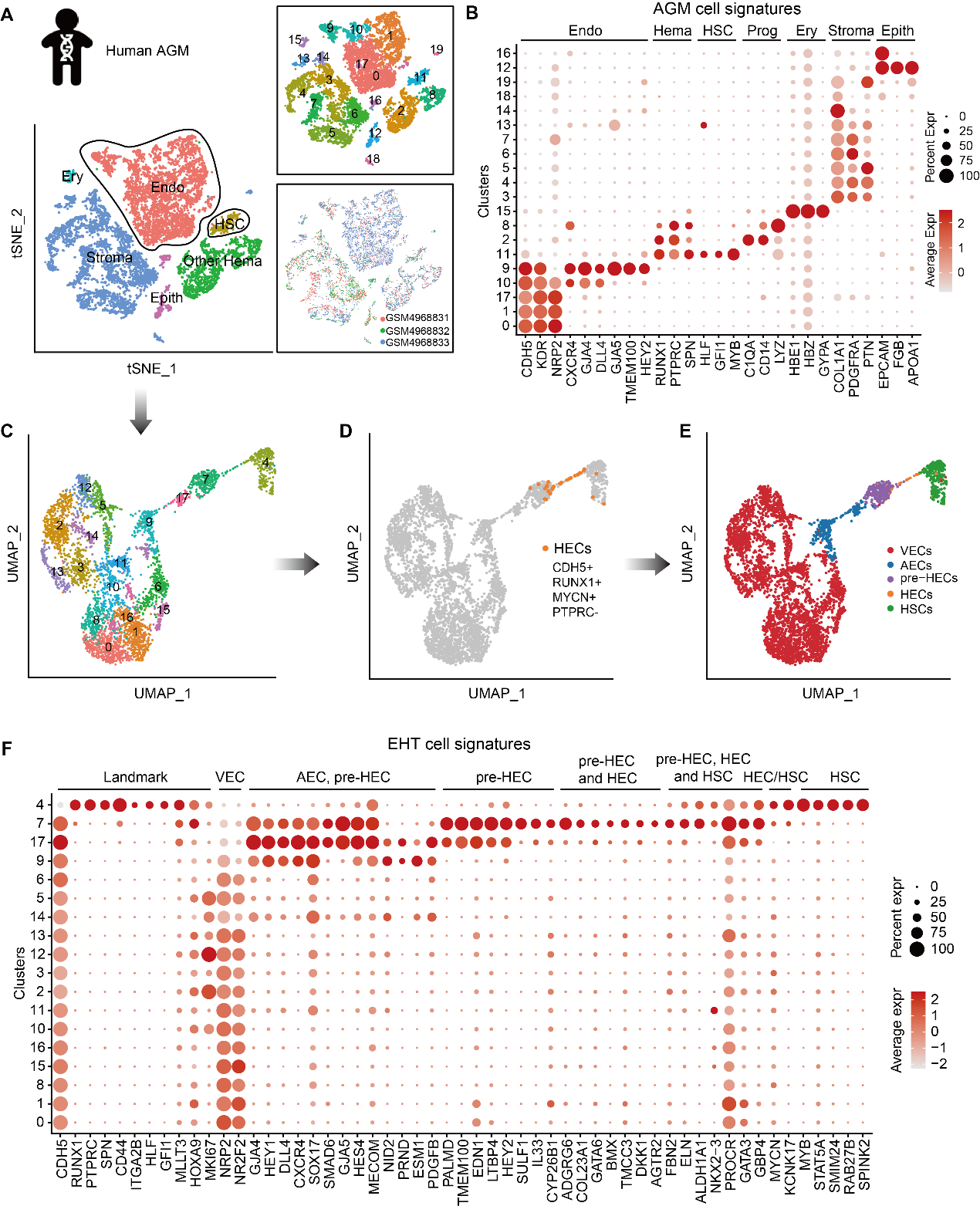

**Figure S1.** Steps to reconstruct the human EHT trajectory. **A** The tSNE plots of human AGM cell types, clusters and samples. **B** Expression of AGM cell signatures in each cluster in (**A**). **C** EHT clusters extracted from (**A**). **D** Annotate HECs by the co-expression of CDH5, RUNX1 and MYCN, along with the absence of PTPRC. **E** The UMAP plot of annotated human EHT cells. **F** Expression of EHT cell signatures in each cluster in (**C**).

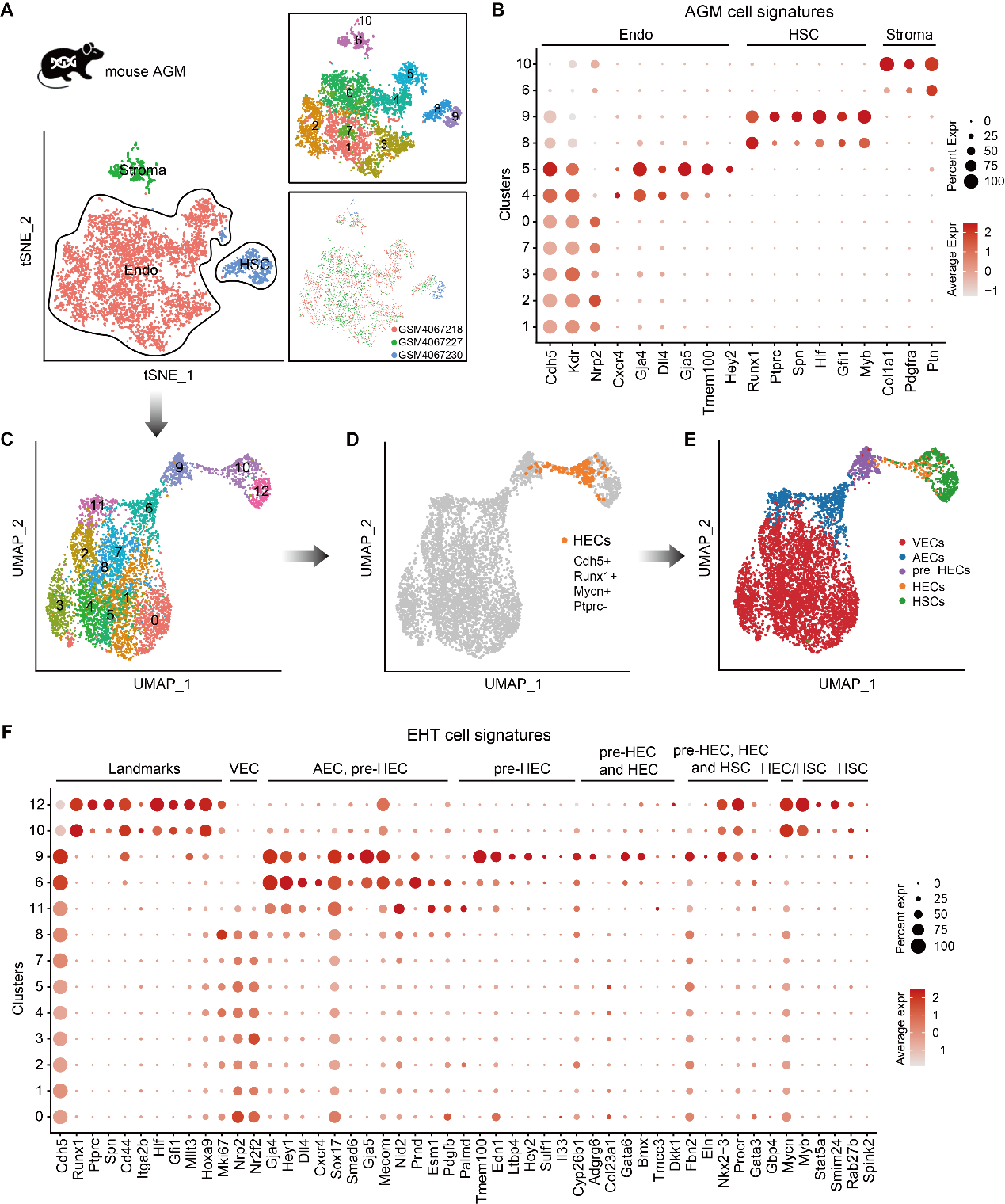

**Figure S2.** Steps to reconstruct the mouse EHT trajectory. **A** The tSNE plots of mouse AGM cell types, clusters and samples. **B** Expression of AGM cell signatures in each cluster in (**A**). **C** EHT clusters extracted from (**A**). **D** Annotate HECs by the co-expression of Cdh5, Runx1 and Mycn, along with the absence of Ptprc. **E** The UMAP plot of annotated mouse EHT cells. **F** Expression of EHT cell signatures in each cluster in (**C**).

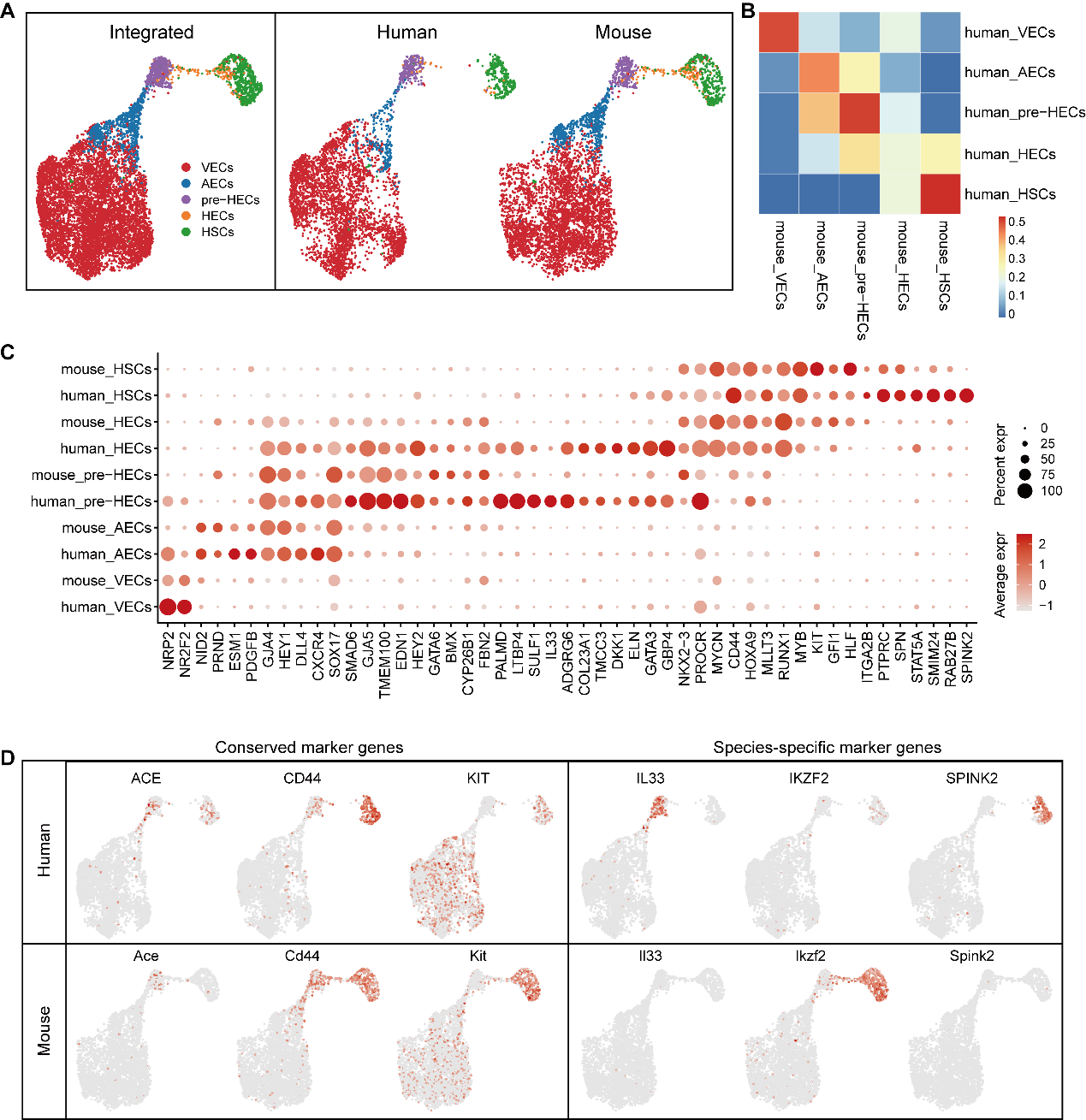

**Figure S3.** Integration of the human and mouse EHT scRNA-seq data. **A** UMAP of integrated human and mouse EHT. The EHT of both showed a highly conserved pattern, although a relatively larger number of HECs were captured in mouse. **B** Correlations of human and mouse EHT cell types. **C** Expression of EHT marker genes in human and mouse EHT cell types. **D** Conserved and species-specific markers between human and mouse EHT.
