## Additional file 9 for "Systematic single-cell analysis reveals dynamic control of transposable element activity orchestrating the endothelial-to-hematopoietic transition"

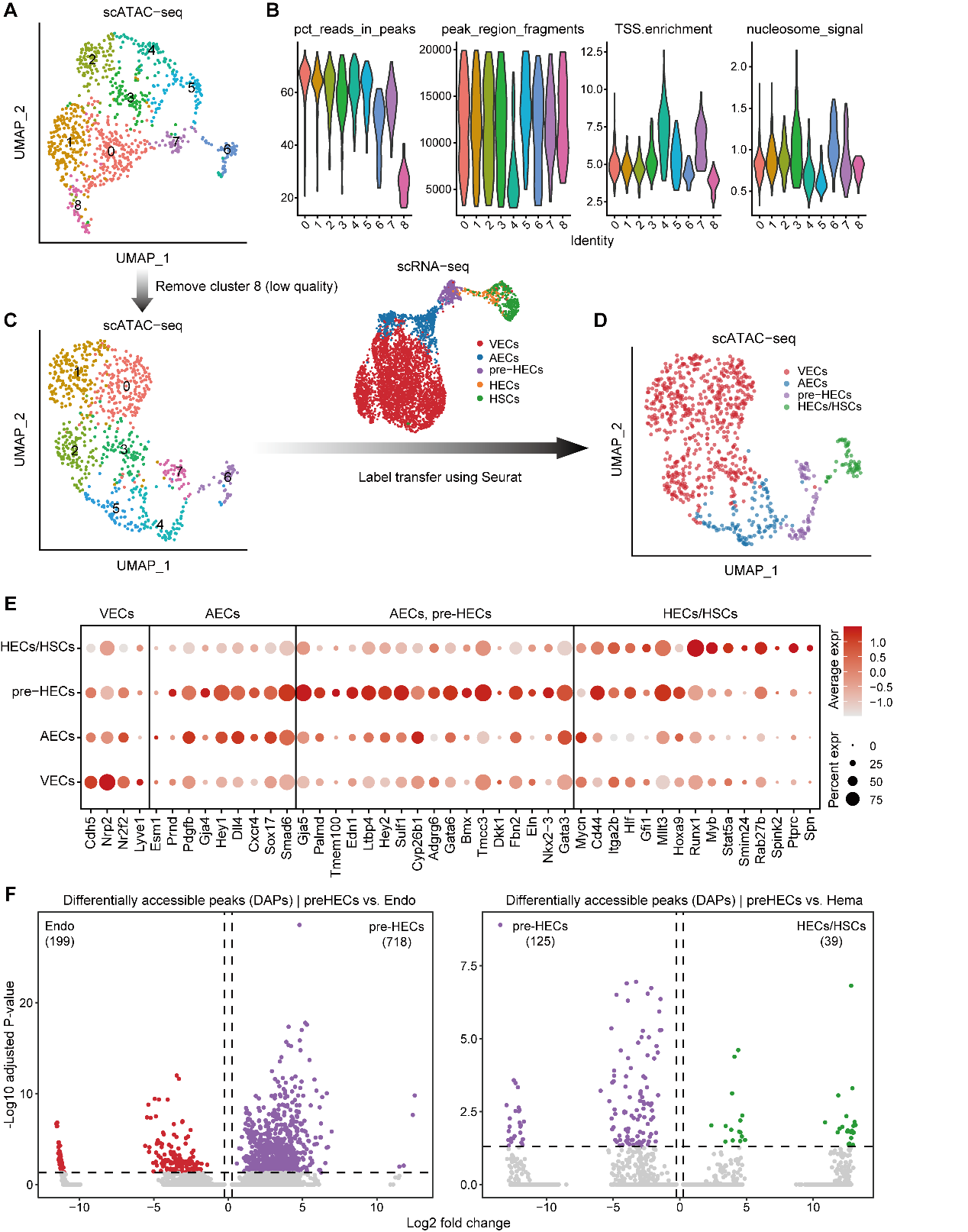

**Figure S1.** Steps to reconstruct the mouse EHT trajectory from scATAC-seq data. **A** The raw UMAP plot of mouse AGM clusters. **B** Quality metrics of mouse scATAC-seq data. **C, D** Filtering and annotating the mouse scATAC-seq data. The cell types are transferred from mouse scRNA-seq data. **E** EHT marker gene activities of mouse EHT cell types in scATAC-seq data. **F** Differential accessible analysis of pre-HECs compared with endothelial cells (VECs/AECs) and hematopoietic cells (HECs/HSCs).

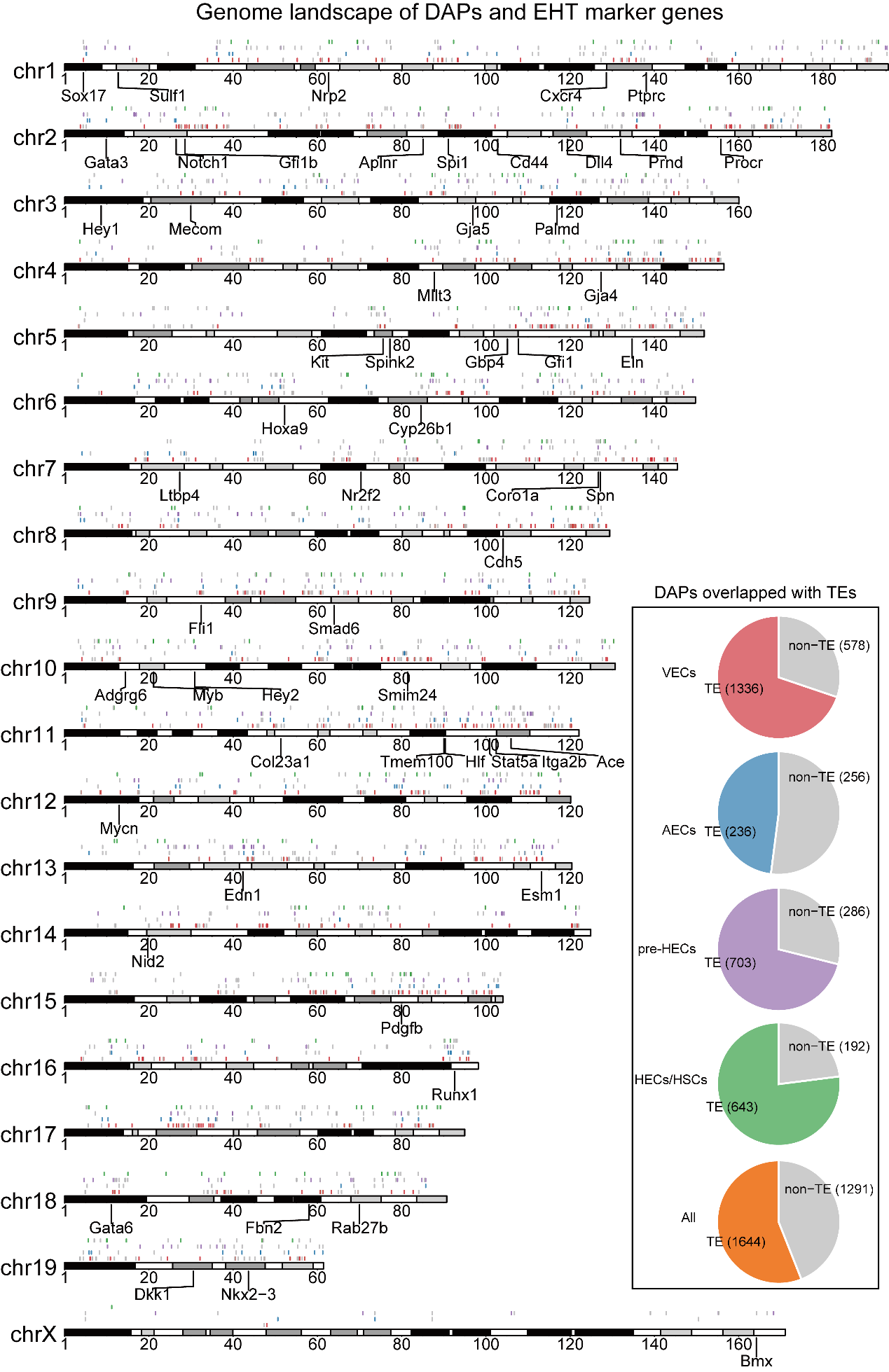

**Figure S2.** Genome landscape of differentially accessible peaks (DAPs). The peaks are grouped into TE and non-TE overlapped. DAPs in different cell types are plotted on four tracks with different colors. Typical EHT marker genes are also indicated on the chromosomes.

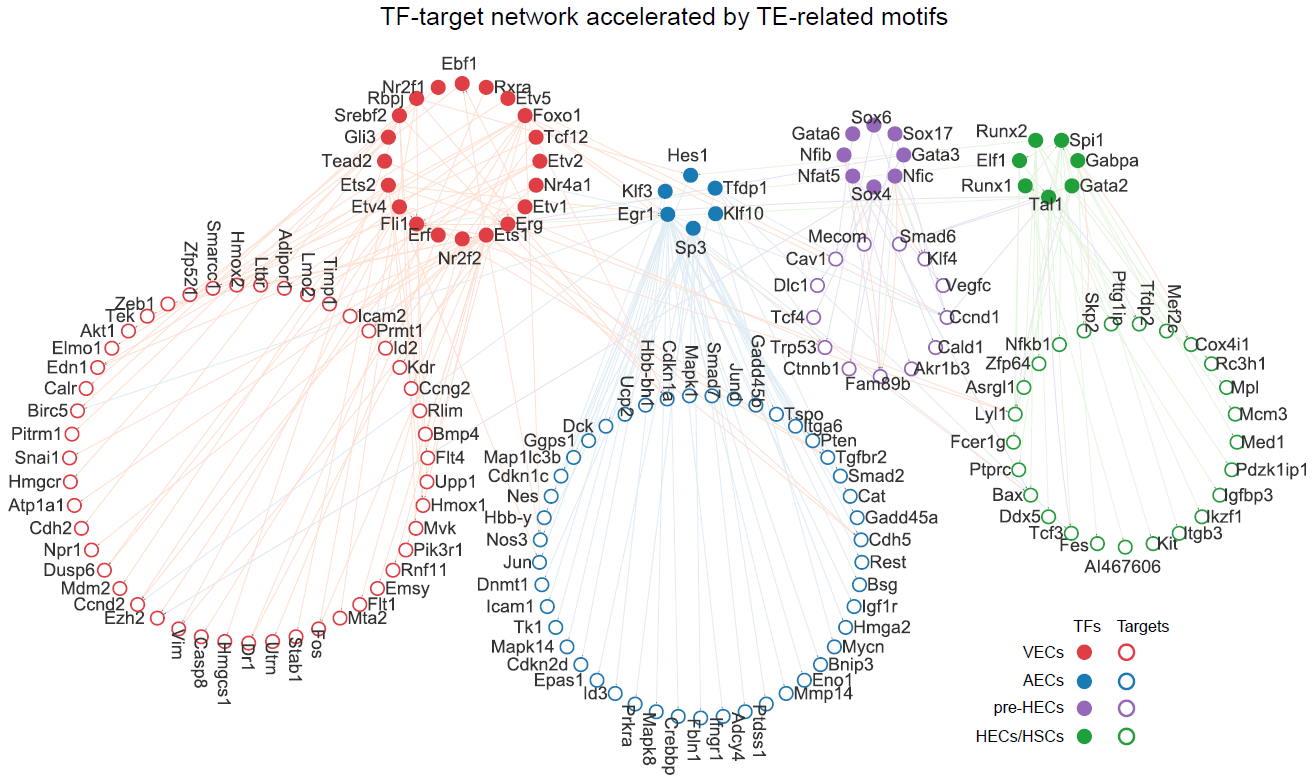

**Figure S3.** The TF-target network in mouse EHT. TFs are selected from TE-related motifs. Targets of TFs are obtained from TRRUST.
