## Additional file 12 for "Systematic single-cell analysis reveals dynamic control of transposable element activity orchestrating the endothelial-to-hematopoietic transition"

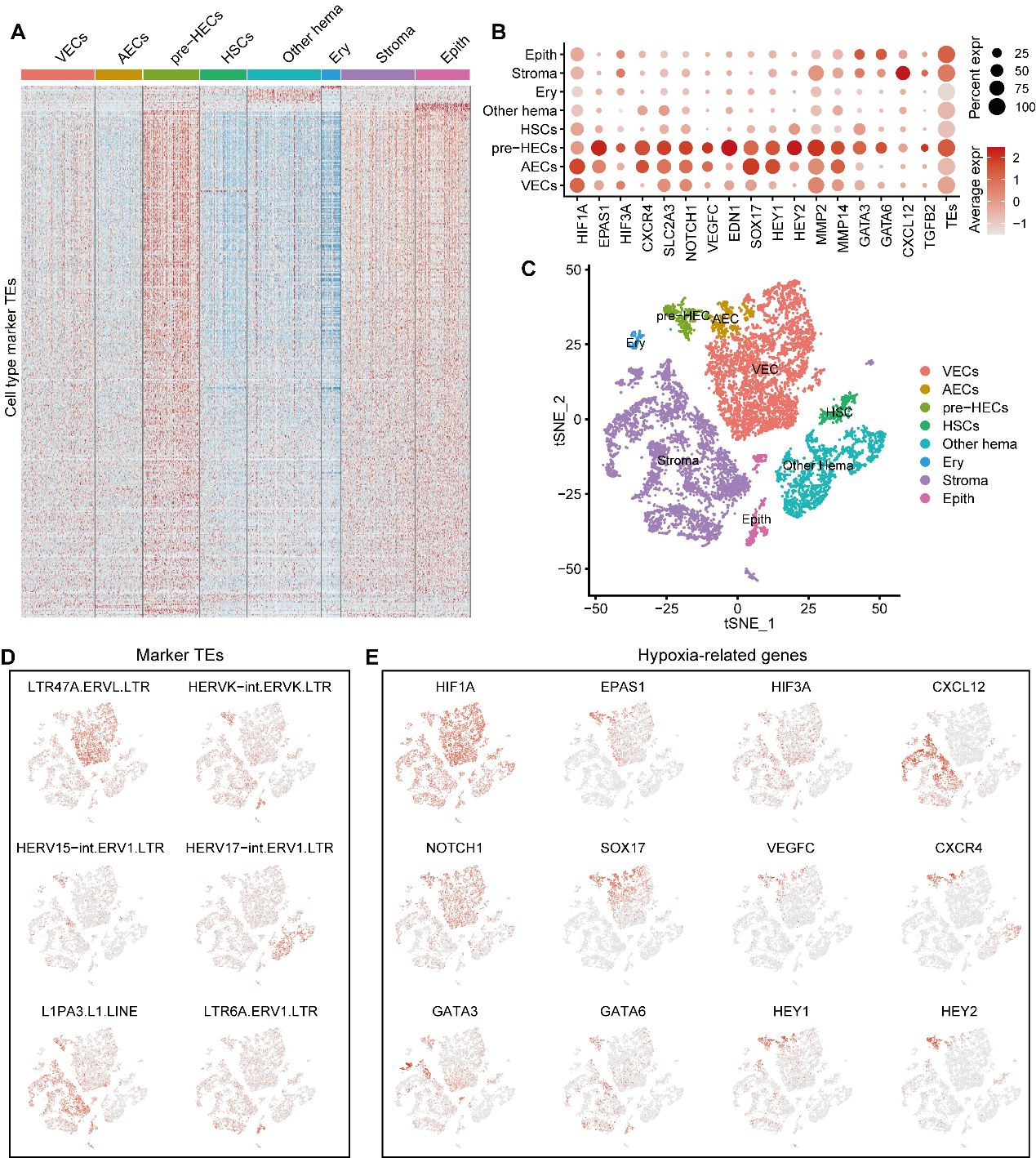

**Figure S1.** The hypoxic niche and TE expression in human AGM scRNA-seq data. **A** Cell type marker TEs in human AGM. In addition to pre-HECs, stromal and epithelial cells also show relatively higher TE expression. **B** Expression of hypoxia-related genes and TE silencers in human AGM. **C** The tSNE plot of human AGM cell types. **D** UMAP of selected marker TE expression in human AGM. **E** UMAP of hypoxia-related gene expression in human AGM.

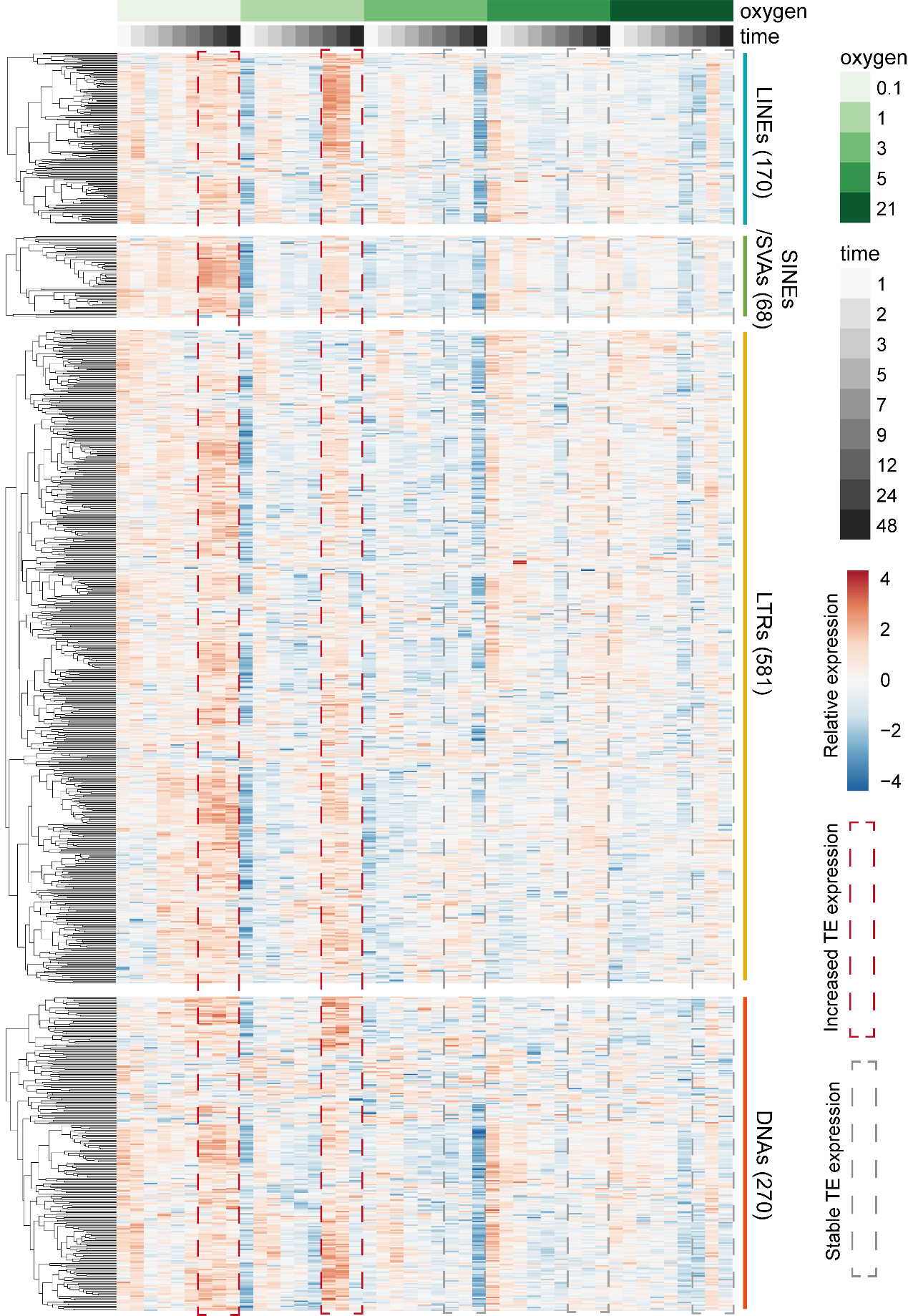

**Figure S2.** Expression heatmap of each TE family (grouped into four TE classes) in HUVEC bulk RNA-seq data.

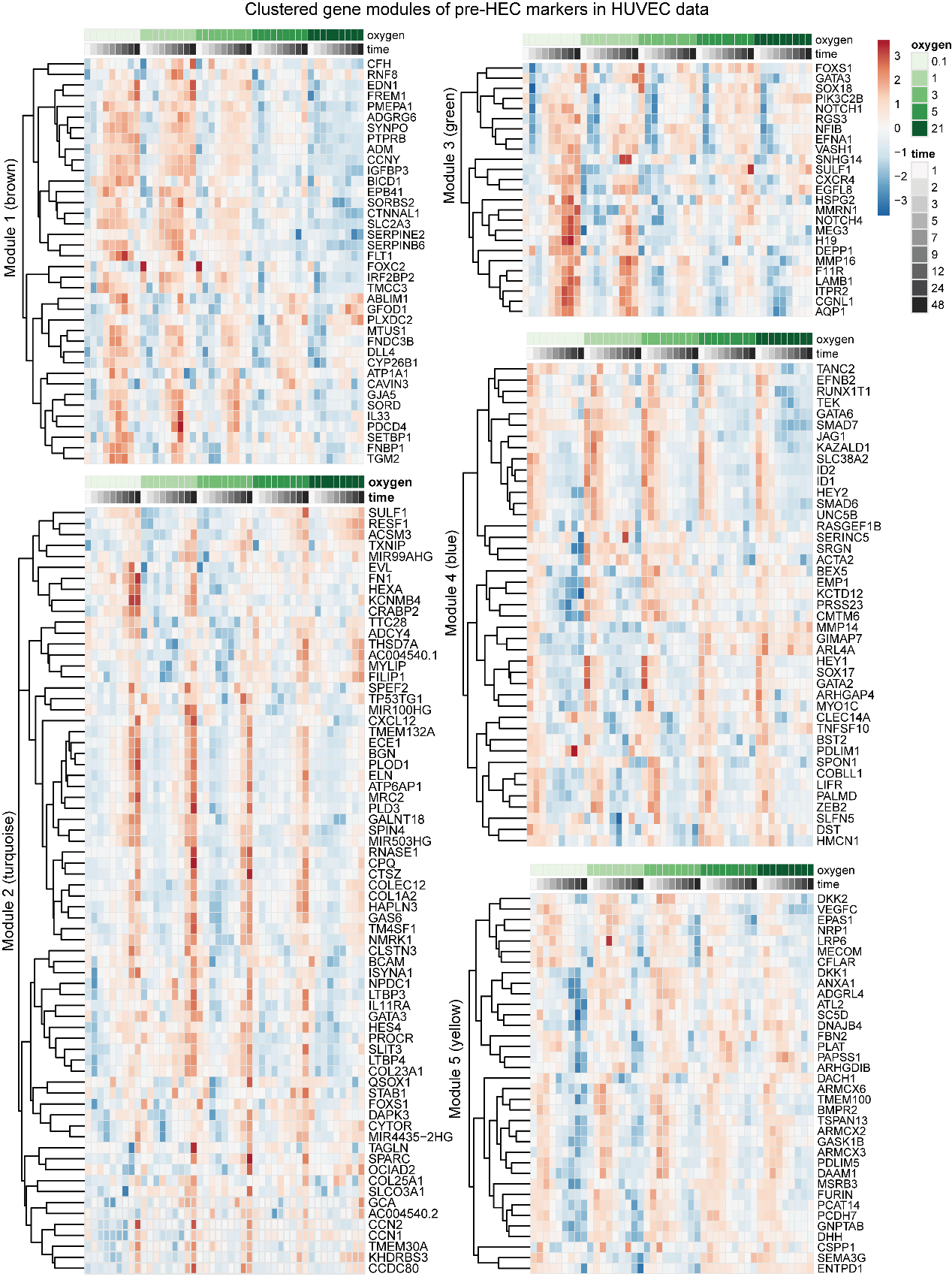

**Figure S3.** Five gene modules of pre-HEC markers on HUVEC data. The gene modules are identified using WGCNA.
